# Influence of pathway topology and functional class on the molecular evolution of human metabolic genes

**DOI:** 10.1101/292714

**Authors:** Ludovica Montanucci, Hafid Laayouni, Begoña Dobón, Kevin L. Keys, Jaume Bertranpetit, Juli Peretó

## Abstract

Metabolic networks comprise thousands of enzymatic reactions functioning in a controlled manner and have been shaped by natural selection. Thanks to the genome data, the footprints of adaptive (positive) selection are detectable, and the strength of purifying selection can be measured. This has made possible to know where, in the metabolic network, adaptive selection has acted and where purifying selection is more or less strong and efficient. We have carried out a comprehensive molecular evolutionary study of all the genes involved in the human metabolism. We investigated the type and strength of the selective pressures that acted on the enzyme-coding genes belonging to metabolic pathways during the divergence of primates and rodents. Then, we related those selective pressures to the functional and topological characteristics of the pathways. We have used DNA sequences of all enzymes (956) of the metabolic pathways comprised in the HumanCyc database, using genome data for humans and five other mammalian species.

We have found that the evolution of metabolic genes is primarily constrained by the layer of the metabolism in which the genes participate: while genes encoding enzymes of the inner core of metabolism are much conserved, those encoding enzymes participating in the outer layer, mediating the interaction with the environment, are evolutionarily less constrained and more plastic, having experienced faster functional evolution. Genes that have been targeted by adaptive selection are endowed by higher out-degree centralities than non-adaptive genes, while genes with high in-degree centralities are under stronger purifying selection. When the position along the pathway is considered, a funnel-like distribution of the strength of the purifying selection is found. Genes at bottom positions are highly preserved by purifying selection, whereas genes at top positions, catalyzing the first steps, are open to evolutionary changes.

These results show how functional and topological characteristics of metabolic pathways contribute to shape the patterns of evolutionary pressures driven by natural selection and how pathway network structure matters in the evolutionary process that shapes the evolution of the system.

## INTRODUCTION

Metabolism is the set of enzymatic reactions that allows the synthesis, degradation and transformation of the biochemical components necessary for the maintenance and reproduction of a cell. Understanding the evolution of a system whose functioning arises from the interplay of many cellular components, is important both, for understanding the biology of the cell and for unraveling general principles of evolution of complex biological systems.

The origin and evolution of metabolic pathways is a difficult problem and several ideas have been proposed (reviewed in Peretó 2011). Among them, the patchwork model has gained a general acceptance. It proposes the evolution of enzymes from broader to narrower substrate specificities through gene duplication and the cooption of metabolic functions by the diverse pathways (Yčas 1974, Jensen 1976). Nevertheless, the ability to contrast different models has been limited by the fact that all of them predate the current availability of complete genome sequences from the three domains of life (Lazcano et al. 1995). Nowadays, complete genome sequences and subsequent reconstructions of genome-scale metabolic networks for many organisms have been used to test some of the predictions of evolutionary models. In the context of those systemic studies, the patchwork model exhibits a higher explicative power (Alves et al 2002; Light and Kraulis 2003; Diaz-Mejia et al 2007; Fani and Fondi 2009; Grassi and Tramontano 2011; Peretó 2012).

A full understanding of the evolution of metabolism also requires the understanding of the functional evolution of the enzymes. This can be achieved by investigating the selective pressures that have been acting on the genes that code for the enzymes (metabolic genes), and trying to understand their evolutionary dynamics in relation to the molecular systems they participate. An interesting approximation is the study of the selective pressures over the network structure of molecular systems; this can be achieved through the study of the relationship between parameters of the evolutionary histories of the enzyme-coding genes and the topological properties of their gene products within a network. Connectivity, which is the number of links of a node in a network, in a metabolic network represents the number of metabolic interactions and it is an initial measure for the topological description of each node.

Selective pressures are at the base of understanding adaptation and two main types have to be distinguished. By one hand purifying selection, which is the force that eliminates genetic variants that impair the function and, by the other, positive selection, in which one (or several) variants show a better adaptation and the frequency in the population will increase, reaching eventually fixation.

These evolutionary pressures may be detected and measured through *dN/dS* when comparing the genomes from different species. A negative correlation between connectivity and the evolutionary rates have been reported in the Drosophila and yeast genome-scale metabolic networks. In these networks, highly connected genes have been shown to evolve at slower rates, indicating selective constraints acting on them (Vitkup et al. 2006; Greenberg et al. 2008). Hence connectivity has an effect on evolutionary rates, with higher connected genes under stronger purifying selection. In mammal genomes, the negative correlation between dN/dS and degree centrality is only found in four sub-cellular compartments while in other three a negative correlation is found between dN/dS and betweenness centrality (Hudson and Conanat 2011).

This same approach, which couples the molecular evolution of genes with the knowledge of the interaction networks of their gene-products, has been also applied to study specific metabolic pathways, and the influence of the local network structure (that is the structure of only the metabolic pathway) is studied. This may reveal local strategies of adaptation that may be found different from the constraints imposed by the whole metabolic network, as well as it may shed light on network constraints specific to only specific pathways.. In these works, the analysis at the network level initially seeks differences in the strength of the selection on genes located upstream versus those located downstream in the pathway, in order to detect whether the position of a gene along the pathway may constrain gene evolution. In plant biosynthetic pathways, it has been found that upstream genes tend to evolve slower than those downstream due to a stronger selective constraint (Rausher et al. 1999; Lu and Rausher 2003; Rausher et al. 2008; Livingstone and Anderson 2009; Ramsay et al. 2009). The analyzed pathways are secondary metabolite biosyntheses, with an inherent directionality given by the consecutive steps of the biosynthetic process. Further, many of the analyzed pathways are organized into branched structures: one or few initial substrates are processed into many final outputs. In such branched pathways with no loops, upstream genes are more likely to be above branch points and hence to be involved in the synthesis of more products than downstream genes (Rausher et al.1999). These branched structures are common to many biosynthetic metabolic pathways. A proposed explanation for the observed gradient in selective pressures is that upstream genes are under stronger purifying selection because they are more pleiotropic than those downstream, affecting a greater number of end products. When the evolution of the genes of the N-glycosylation pathway during the divergence of primates was analyzed, an opposite trend emerged, with genes at the downstream position of this metabolic pathway being more constrained than upstream ones (Montanucci et al 2011). In a non-metabolic pathway, phototransduction, proteins which are topologically central in the signaling pathway, were found to be more constrained in their evolution; proteins peripheral to the pathway have experienced a relaxation of selective pressures (Invergo et al 2013).

All these studies suggest the idea that a part of the variation of evolutionary rates in metabolic pathways can be accounted for by the structure of their functional network, both the local metabolic pathway and the global whole-cell metabolic network. However, in the case of single metabolic pathways, different patterns have been found for different pathways and different species sets and no general pattern has emerged from these few cases. A large number of pathways and an overall vision of the metabolic network should be analyzed to reveal whether there exist general patterns in their evolutionary history and to better understand the distribution of selective pressures in the network, both in terms of constraints or adaptations. Here, we address the relationship between the topology of the whole set of human metabolic pathways during the time of divergence of primate and rodents and relate it to the evolutionary behavior of each gene in terms of natural selection (purifying and adaptive). Its relationship may help understand the distribution of evolutionary forces within complex biomolecular networks.

## METHODS

### Data Set

#### Pathways

The data set is composed of the metabolic pathways comprised in the HumanCyc database, release 18.1. The number of human pathways present in HumanCyc 18.1 is 325. Of these, 13 pathways were not considered in the present study because they are signaling or protein modification pathways. Two pathways (morphine biosynthesis and melatonin degradation III) could not be used because their reactions are not associated to any annotated human gene. The final total number of considered metabolic pathways for the analysis is then 310. Of these, 275 are base pathways (comprised of reactions only) while 35 are super-pathways, which are comprised of one or more pathways, plus possible additional reactions. These super-pathways were treated separately in the analysis because they add redundant information. A full list of the pathways can be found in Supplementary Table S1.

The election of HumanCyc may requires justification. Even if there are other sources of reconstruction of human metabolism (Duarte et al. 2007; Ma et al. 2007; Thiele et al. 2013), in our case the choice of the HumanCyc database was precisely driven by the fact that it annotates pathways, and the study of the influence of the local topology (pathway topology) on the evolution of metabolic genes was precisely the question motivating this work. Our main objective was to gather the best possible functional, expert-curated annotations within a pathway classification to be further manually curated and our examination indicated that HumanCyc had the best quality collection of pathways.

#### Reactions

The total number of enzymatic reactions comprised in the 310 pathways that are associated to at least one annotated gene is 879.

#### Genes

The total number of genes that encodes enzymes that take part in the annotated pathways is 956. Genes were functionally classified by assigning them to the functional class of the pathway in which they participate. Two different classification schemes were considered since there are different criteria according to which pathways can be assigned to distinct functional groups. The first classification scheme used, the ontology-based scheme, relies on the HumanCyc Pathway-ontology that aims at classifying pathways into a tree-based structure (as a gene ontology). We used the top level of the tree (the classes immediately below the root of the tree, which is “Pathway”) as classes to assign pathways. We used 7 of the 9 parent classes just after the root of the HumanCyc pathways ontology, excluding the two classes for non-metabolic pathways, “Macromolecule Modifications” and “Signal transduction pathways”. The basic criteria of this ontology rely on the metabolic mode of the pathway, for example biosynthesis versus degradation. The second classification schema, compound-based, is based on the kind of compounds that are primarily transformed in the pathway, for example nucleotide versus fatty acids (see classification in supplementary Table S1).

### Computation of Evolutionary Rates

Evolutionary rates were estimated during the divergence of mammals and rodents. For each human gene, its orthologous sequences were derived from the following species: chimpanzee, orangutan, gorilla, mouse and rat. Multiple sequence alignments of the coding regions were downloaded from Ensembl (release 75). When absent, orthologous sequences were predicted, if possible, through a similarity search of the human gene sequence against the genome assembly, followed by subsequent gene prediction by GeneWise (Birney et al 2004), in a procedure described in Montanucci et al (2011). In case of predicted orthologues, multiple sequence alignments were obtained through T-coffee with default options by aligning protein sequences and then back-translating to genomic sequence. The same procedure was adopted for incomplete or bad quality sequences.

Evolutionary rates were computed using the codeml program of the PAML package (Yang 2007). Two likelihood ratio tests between pairs of nested model (M1a versus M2a and M7 versus M8) were carried out to detect positive selection events. The overall strength of purifying selection on each gene was estimated though a unique *dN/dS* over the entire tree and sequence length (model M0). Each maximum-likelihood estimation, including likelihood ratio tests, was carried out 5 times, each with 3 different initial *dN/dS* values: 0.1, 1 and 2 to check for stability of the results to repeated runs and different initial conditions. Final results of the likelihood ratio tests were corrected through a False Discovery Rate (FDR) method (Storey 2002). Positive selection was inferred when either one or both of the two likelihood ratio tests was significant after correcting for multiple testing. Given the shallow divergence considered, non-branch-specific models (M0 averaged dN/dS and site-specific positive selection tests) provide the best estimation of the overall selective pressure acting on each gene, given that for low number of closely related species branch-specific estimations that lack the power to provide stable estimations.

Given the strong impact of alignment errors in generating spurious signals of positive selection, the alignments corresponding to genes with P < 0.05 in the likelihood ratio test for positive selection were inspected visually. Alignment regions containing evident errors (usually contiguous positions in a sequence of the alignment with a suspiciously high number of differences often resulting from sequencing or assembly errors) were manually masked and then evolutionary rates and positive selection tests were then recomputed.

### Computation of Topological Parameters

#### Building the reaction graph

Metabolic pathways were derived from HumanCyc. The classical representation of metabolic pathways, also used in HumanCyc, is through the substrate graph, in which nodes represent metabolites and edges represent reactions that transform metabolites. Here we represent pathways as reaction graphs (Montañez et al 2010), in which nodes represent reactions and edges link consecutive reactions. Consecutive reactions within a pathway were derived from the PREDECESSORS field of the pathways.dat flat-file of the HumanCyc database. The direction provided in this field was used to build directed graphs, in which edges are arrows having a direction that goes from the preceding to the following reaction. The graphs have been constructed through Python scripts from the flat-files downloaded from HumanCyc 18.1.

#### Centrality measures

Topological measures within the directed graphs built for each pathway independently were computed through build-in functions of the NetworkX Python package. Four centrality measures were computed: closeness, betweenness, and in-degree and out-degree centrality. Centrality measures for the corresponding undirected graphs were also computed.

#### Loops

A Boolean variable was derived for each pathway to detect whether the pathway contains loops. This computation has been achieved through the *simple_cycles()* function of the NetworkX package applied to the directed graph. This variable was only computed for pathways of more than one reaction.

#### Top/Bottom Position

For each pathway, nodes were assigned to three different classes depending whether they are at the beginning of the pathway (in-degree=0, “upstream”), at the end (out-degree=0, “downstream”) or in any other position in between (“intermediate”). The uppermost positions (top positions), are the first reaction steps of the pathway, corresponding to initial reactions. At bottom positions there are the reactions that produce the final products of the pathway. Reactions in any other position in between of these two are assigned to the same class (intermediate). Beside these three classes, a fourth class has been added for nodes (reactions) that become isolated when the substrate graph is transformed into the corresponding reaction graph. This is the case for pathways for which the reaction graph comprises more than one connected component. These reactions have the feature of directly catalyzing the end products of the pathway starting from the initial substrates, thus being at the beginning (first step) and at the end (last step) at the same time.

### Statistical Analyses

To evaluate the importance of relationship between evolutionary estimates and the different descriptive network properties, we performed a multivariate analysis, through an automated linear modeling routine implemented in SPSS software. The automated linear modeling created a single standard model to explain the relationship between fields. The adopted linear modeling also included codon bias, protein length and CG content as explanatory variables. Before the linear modeling missing values were replaced. No variable selection method was used and all variables entered into the model were assessed. The automated linear modeling allows also for detecting outliers; however outliers were not excluded of the analysis. To compare groups, non-parametric methods were used. To correct for multiple testing, False Discovery Rate (FDR) methods were applied (Storey 2002).

Genes considered under positive selection were compared to the whole set of metabolic genes for several centrality measures using a permutation test. For every centrality measure, the mean score of the genes under positive selection was compared to the distribution of the mean scores of a set of randomly selected genes from the whole set of metabolic genes with the same sample size, using a permutation test (10.000 permutations).

## RESULTS

### Data Set Description

The total number of pathways used for the analysis is 310, with 275 base pathways and 35 super pathways (see methods). A full list of the pathways can be found in supplementary Table S1. Of the 275 base pathways, 30 comprise only one reaction, while the remaining 245 pathways comprise a number of reactions that ranges from 2 to 30. The majority of the pathways (208 out of 275 base pathways and 28 out of 35 super pathways) show no loop structures. They are characterized by a non-feedback topology, with either linear or branched structures. For 27 of the 35 super-pathways, their reaction set is fully comprised in base pathways, so excluding them from the analysis would not result in loss of reactions and genes.

The total number of genes associated to the reactions in the pathways is 956. 943 out of the 956 gens participate in base pathways, while 13 genes (*NAGK*, *GGCT*, *SRR*, *TYW4*, *GGT1*, *EBP*, *SC5DL*, *DHCR7*, *DHCR24*, *CCBL1*, *OPLAH*, *CNDP2*, *CCBL2*) are associated to reactions that are uniquely present in super pathways. Of the 956 genes, 335 encode proteins that carry out their enzymatic activity within protein complexes while 621 genes encode proteins that are themselves the functional enzymes.

The relationship between enzymatic activities and enzymes is not one-to-one: on the one hand, each gene encodes an enzyme (or an enzymatic subunit) that may carry out more than one catalytic activity and, on the other hand, the same catalytic activity can be served by more than one enzyme (isoenzymes) that are encoded by different genes. Within this scenario, 71% of the genes (677 out of 956) code for enzymes (or enzymatic subunits) that carry out only one metabolic function, 26% of the genes (247 out of 956) are associate to a number of reactions between 2 and 5, and the remaining 3% of the genes (32 out of 956) are associated to more than 5 reactions. The genes whose encoded enzymes are associated to the biggest number of reactions are: *FASN* (fatty acid synthase), *CYP2A6* (a type of cytochrome P450), *UGT2B11* (a type of UDP-glucuronosyltransferase), *SCMOL* (a methylsterol monooxygenase) *DIO3* (deiodinase, iodothyronine, type III), *ALOX5* (arachidonate 5-lipoxygenase) and *ALDH3A2* (fatty aldehyde dehydrogenase).

### Purifying and Positive Selection in Metabolic Genes

The evolutionary rates have been computed for 927 genes (for 29 it was not possible due to lack of or poor quality of one or more of the orthologous sequences) considering the full set of six species (human, chimpanzee, orangutan, gorilla, mouse and rat): the synonymous evolutionary rate (*dS*), the non-synonymous evolutionary rate (*dN*) and their ratio (*dN/dS*), which provides the direction of the action of the selection and an overall estimation of the strength of the purifying selection that have acted on each gene. The distribution of the evolutionary rates can be seen in Fig 1, which shows that the main force that has shaped the evolution of metabolic genes during the mammal evolution has been purifying selection, with all *dN/dS* values lower than 0.5.

**Figure 1:**
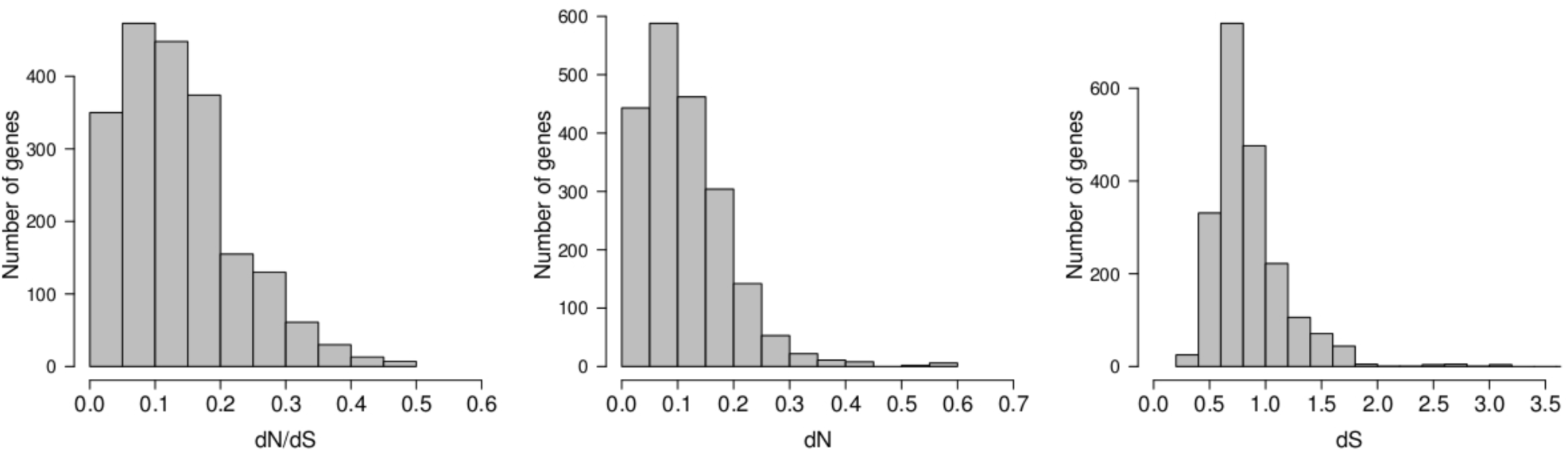
Distribution of the evolutionary rates (*dN/dS*, *dN* and *dS*) for the 927 metabolic genes.

Beside an estimation of purifying selection, for each one of the 927 genes we performed two likelihood ratio tests to look for sequence signature of positive selection. After multiple test correction, using false discovery rate, we found that only six genes show a significant P-value for the likelihood ratio test. The genes and their p values for the M7 versus M8 test are: CYP2E1 (P = 0.0000005, corrected = 0.00043), HDC (P = 0.0000217, corrected = 0.01006), CES1 (P = 0.0000471, corrected = 0.01455), DPM2 (P = 0.0001017, corrected = 0.02356), SPAM1 (P = 0.0001575, corrected =0.02919) and AKR1C1 (P = 0.0002451, corrected =0.03787).

CYP2E1 is a member of the cytochrome 450 family of enzymes involved in the inactivation of drugs and xenobiotics. The HDC gene codes for histidine decarboxylase, which converts L-histidine into histamine, a biogenic amine involved in different physiological processes such as neurotransmission, gastric acid secretion, inflammation and regulation of circadian rhythm. CES1 codes for a carboxylesterase involved in the hydrolysis of various xenobiotics and drug clearance in liver. DPM2 encodes for the regulatory subunit of the dolichol-phosphate mannose (DPM) synthase complex whose main function is recognition on the cellular surface. SPAM1encodes for a hyaluronidase located on the human sperm surface that enables sperm to penetrate through the hyaluronic acid-rich envelope of the oocyte. AKR1C1 belongs to a superfamily of aldo-keto reductases and catalyzes the reaction of progesterone to the inactive form 20-alpha-hydroxy-progesterone.

The quality control of the alignments which involve the removal of bad quality regions from the computation of the evolutionary rates might have led to an underestimation of the positive selection events, however it guarantees the reliability of the events found.

### Functional classes

Beside the evolution of specific genes under positive selection, we investigated different levels of selective constraint between functional classes by comparing evolutionary rates of genes participating in pathways performing different functions. A Kruskal-Wallis (KW) test was used to test differences in evolutionary rates between genes belonging to different functional classes of pathways. For both ontology-based and compound-based classification, we find significant differences among different functional classes in the evolutionary rates *dN*, and *dN/dS* (with *dS* close to significance).

When we consider the ontology-based classification (supplementary Fig S1), differences in *dN/dS* reveals relaxed constraints (high *dN/dS* values) for external routes (“Detoxification”), and strong constraints (low *dN/dS*), for routes of the core metabolism (“Generation of precursor metabolites”). We also find that biosynthesis routes are more constrained than degradation ones. These differences in the strength of purifying selection can reflect the need of making the biosynthesis of very specific metabolites, but degradation of a broad range of external compounds, most likely unknown and toxic.

When we consider the compound-based classification (supplementary Fig S2), values of *dN/dS* separate those with highest values, Steroid, Secondary metabolism and Detoxification, and those with the lowest, mainly Glycolysis/TCA/PentoseP, which corresponds to the inner core of the metabolic network. In particular, the detoxification class, the one with the highest *dN/dS*, contains pathways with enzymes able to recognize a broad range of metabolites. Thus, both classification schemes show that genes participating in peripheral routes have evolved under relaxed constraint, while stronger selective constraint has acted on the genes whose encoded enzymes have roles within the central metabolism.

From a conceptual point of view, traditionally metabolism has been regarded as a series of layers of complexity, from central pathways involved in basic energy transactions and the generation of intermediate precursors, followed by a tier of almost universal biosynthetic pathways that use a few of those precursors to generate biomass components, and, finally, some processes connected to central intermediates that generate a diversity of metabolites usually related to behavioral or environmental cues, and showing a more restricted phylogenetic distribution, also known as secondary metabolism. Theoretical approaches based on graph theory support this classical image of metabolic organization (Guimerà and Amaral 2005; Noor et al 2010). Thus, we grouped the functional classes into three main groups according to the layer of the global metabolism in which they operate: the inner core of intermediary metabolism comprises classes of Glycolysis/TCA/PentoseP and Polysaccharides; the second layer of intermediary metabolism with Membrane Lipids, Nucleotide, Fatty acid/TAG, Cofactor, Fatty Acid/hormone, and Amino acid; and the outer layer of metabolism comprises the classes of Steroid, Secondary Metabolism and Detoxification. When we compare the evolutionary rates of these classes through a KW test we find highly significant differences with a clear gradient of selective constraints in which central routes of the core metabolism are more constrained (lower *dN* and *dN/dS*) than the ones in the second layer which, in turn, are more constrained than those of the more peripheral layer (Fig 2 and supplementary Fig 3). Pairwise comparisons hold significant after correcting for multiple testing (P < 0.001). Figure 2 clearly shows the steep gradient of selective pressures with genes of the central metabolism under the stronger selective constraint and genes of the peripheral routes under relaxed selective constraint.

**Figure 2:**
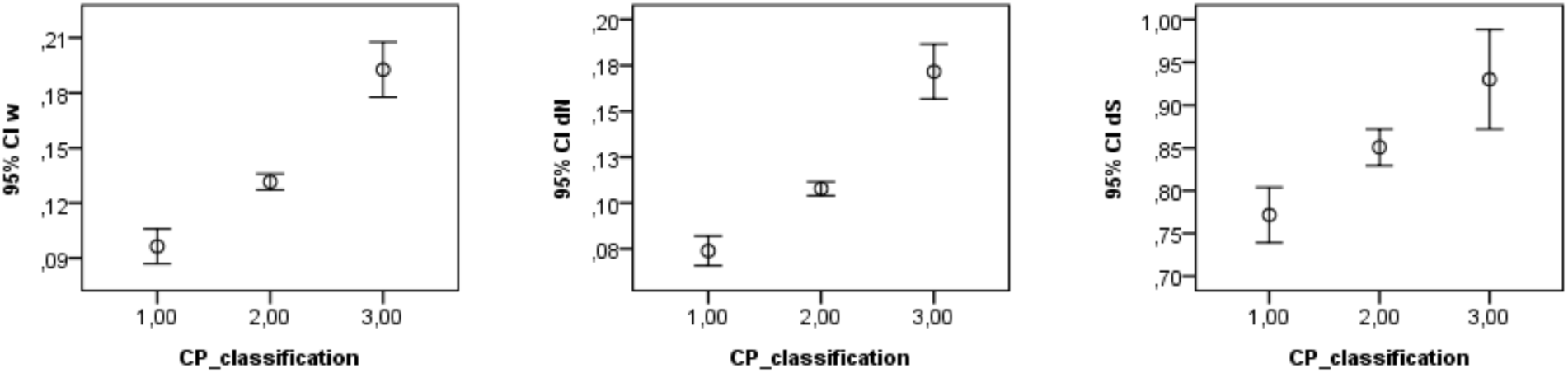
Graph representing *dN/dS*, *dN* and *dS* among genes belonging to different functional classes. Class 1 comprises the inner metabolism (Glycosis/TCA/PentoseP, Polysaccharides), class 2 comprises the second layer (Membrane Lipids metabolism, nucleotide metabolism, Fatty acid/TAG, Cofactor, Fatty Acid/hormone, and Aminoacid) while class 3 comprises the outer layer of cell metabolism (Steroid, Secondary Metabolism and Detoxification).

**Figure 3:**
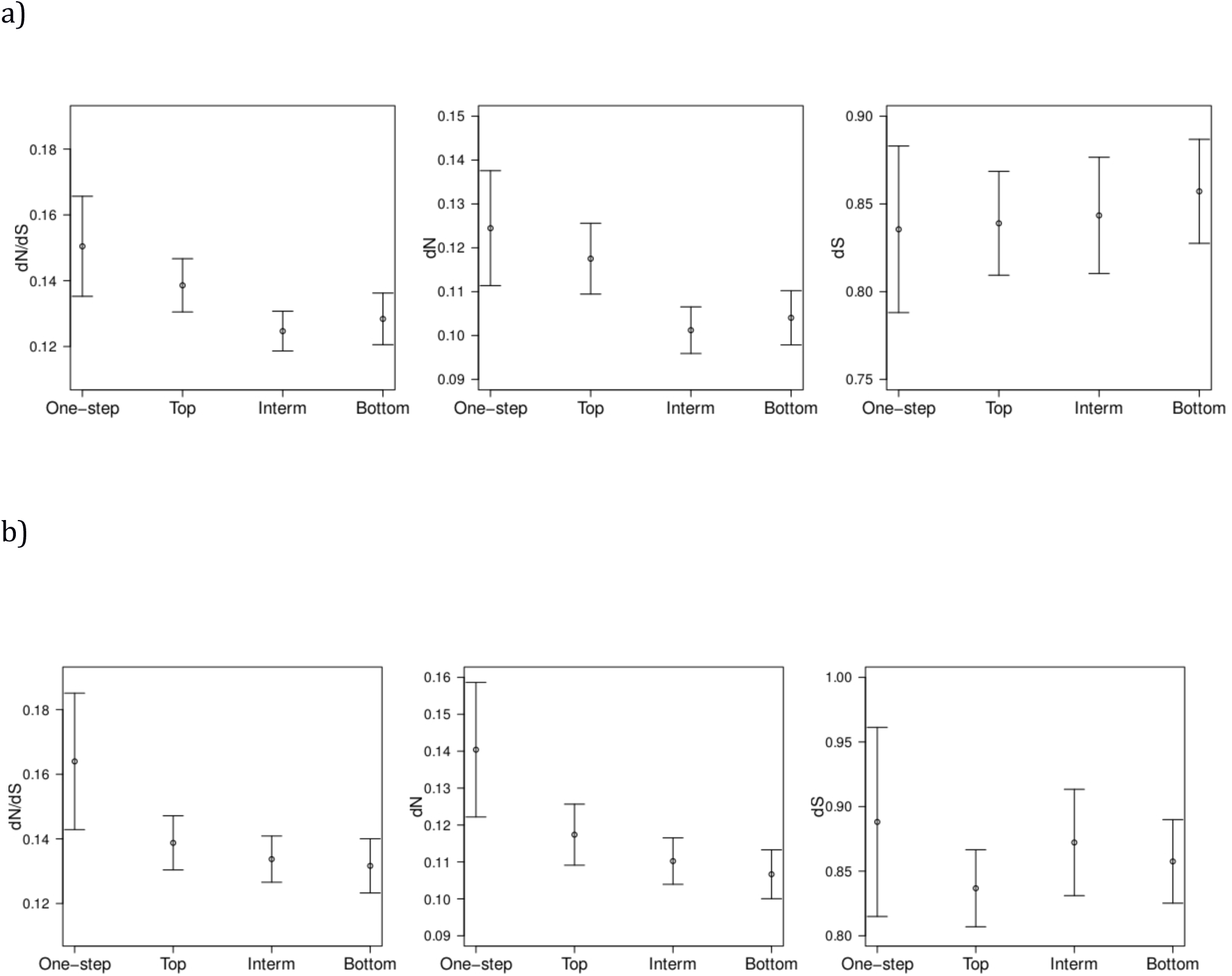
**a)** Graph representing *dN/dS*, *dN* and *dS* among genes whose position within the pathway belongs to four classes (one-step, top, intermediate, bottom positions) for the 275 base pathways. **b)** The same for the 208 base pathways with no loops. Dots show the mean ± 2 standard error (SE).

### Presence/Absence of Isoenzymes

Given that roughly 36% of the reactions are catalyzed by more than one enzyme (different enzymes, encoded by different genes, able to catalyze the same reaction or isoenzymes) we compared the selective pressures that have acted on genes that encode for unique enzymes, to that of genes that encode for enzymes for which isoenzymes exist. The existence of isoenzymes indicates that alternative proteins could be recruited to perform the same metabolic function, whereas the presence of a unique enzyme is critical for their specific metabolic function to be served. Interestingly, we find no difference in selective constraint between these two classes of genes. A Mann-Whitney U test shows no significant differences: P=0.066 for *dN/dS*, P= 0.109 for *dN* and P= 0.668 for *dS*. This result shows that sequence properties of isoenzymes do not differ from those of the rest of the enzymes; they cannot be considered simply as redundant.

### Evolutionary Rates and Topological Properties

In order to investigate whether and how the organization into metabolic networks imposes constraints on the evolution of the enzyme-coding genes, we carried out automated linear modeling to reveal possible linear relationships between evolutionary rates and topological features of reactions (Table 1). Among the topological features that can be associated to each node and that summarize aspects of the node’s position within the network, we first considered four centrality measures: (i) in-degree centrality, which is indicative of the number of incoming links pointing to a node; (ii) out-degree centrality, which is indicative of the number of outgoing links stemming from a node; (iii) betweenness centrality, which is indicative of the importance of a node in linking parts of the networks; and (iv) closeness centrality, which is indicative of whether a node lies in the central or peripheral part of the network. In this analysis, pathways with the category “one-reaction” are excluded, given that topological measures by definition cannot be computed. In the multivariate statistical analysis we included three sequence features of the genes that are known to influence evolutionary rates and thus have to be taken into account when analyzing variations in evolutionary rates. These three sequence features are: (i) ENC, the effective number of codons, which quantifies codon bias and is correlated with expression levels; (ii) length of the coding sequence (CDS) of the gene; and (iii) CG content of the CDS.

**Table 1.** Results for the three independent runs of automated linear modeling with evolutionary measures as dependent variable for the 275 base pathways. The following variables have been considered by the model: 3 gene sequence properties, effective number of codons (ENC), gene length (Length) and CG content (CG); and 4 topological centralities: closeness, betweenness and in-degree and out-degree centralities. Beta coefficients and P-values (between brackets) are reported; significant values according to a threshold of 0.05 are presented in bold.

|  | ENC | Length | CG | Closeness | Betweenness | In-degree | Out-degree |
| --- | --- | --- | --- | --- | --- | --- | --- |
| *dN/dS* | **0.003**<br>(**<0.0001**) | **-0.000**<br>(**<0.0001**) | 0.002<br>(0.958) | -0.056<br>(0.071) | -0.017<br>(0.225) | **-0.035**<br>(**<0.0001**) | 0.038<br>(0.246) |
| *dN* | 0.000<br>(0.901) | **-0.000**<br>(**<0.0001**) | 0.022<br>(0.529) | -0.019<br>(0.502) | -0.021<br>(0.111) | **-0.030**<br>(**<0.0001**) | 0.011<br>(0.703) |
| *dS* | **-0.025**<br>(**<0.0001**) | **-0.000**<br>(**<0.0001**) | **0.549**<br>(**<0.0001**) | 0.021<br>(0.866) | 0.054<br>(0.339) | -0.019<br>(0.573) | -0.037<br>(0.779) |

As seen in Table 1, synonymous substitution rates show significant linear relationships only with sequence features (codon bias, sequence length and CG content), while no significant correlation is found with any of the topological parameters. This implies that, as expected, neutral evolution (evolution at synonymous sites) is only affected by the nucleotide composition of the sequence itself and not by the position of the gene in the network. On the contrary, in the case of functional evolution (evolution at non-synonymous sites and *dN/dS*) a significant linear relationship is found with topological parameters, showing that the evolutionary rates suffer the influence of the position and the role of gene products within the pathways. In particular a highly significant negative correlation is found between both *dN* and *dN/dS* and in-degree centrality, meaning that genes that have high in-degree centralities are highly constrained in their evolution with a strong purifying selection, while genes that have low in-degree centralities are free to evolve under relaxed constraints and accumulate non-synonymous substitutions at a faster rate. This result holds for pathways of different topologies regardless of whether the structure presents loops or not.

### Position along the Pathway: Top/Intermediate/Bottom Classification

In order to analyze the possible relationship of topological centrality measures and the position within the specific metabolic pathways, we took advantage of the physiological directionality of metabolic pathways and we generated a new categorical variable to account for position. Three different classes of reactions have been defined according to their position within the pathway: reactions that catalyze the first enzymatic step in the pathway (top position); reactions that catalyze the last step in the pathway (bottom position); all the in-between steps are assigned to same class (intermediate). A fourth class has been introduced for reactions that catalyze the first and last step at the same time, being, for example the only reaction of one branch of a branched pathway that directly converts the initial substrates into the final products. Genes are assigned to the class of the reaction catalyzed by the enzyme they encode. It should be noted that this classification on position is correlated with in-degree centrality: nodes at top positions are those with no incoming links (in-degree equal to 0) and those at bottom positions are defined solely on the basis of out-degree (out-degree equal to 0). This new variable, even if correlated with in-degree, encodes different positional information that is not fully captured by the in-degree centrality.

We analyzed whether genes whose encoded enzymes catalyze reactions belonging to these four classes have evolved under different selective pressures. Evolutionary rates for these four classes are shown in Fig 3a and supplementary Fig 4a. The statistical significances of their differences have been assessed through Kruskal-Wallis, which show that there are significant differences between the four considered classes in *dN/dS* (P=0.001) and *dN* (P=0.001) while no significant differences are found in *dS* (P=0.151). Both cases of *dN/dS* and *dN* pairwise comparisons show that the intermediate class has a lower *dN* and *dN/dS* than the top (0.022 for *dN/dS* and 0.010 for *dN*) and one-step (0.010 for *dN/dS* and 0.005 for *dN*) classes. This means that genes whose encoded enzymes catalyze reactions at top positions have undergone more relaxed evolution and faster non-synonymous substitution rates than genes at intermediate position. Genes belonging to the one-step class have the highest evolutionary rates.

Given that the last three classes (top, intermediate, and bottom) constitute an ordered variable of the position along the pathway, we carried out a trend test to test whether a linear relationship exists of the evolutionary rates and these positional classes. A significant linear relationship is found for *dN* for the three classes of top-intermediate-bottom with P-value of 0.008. Non-significant P-values for the linear relationship are instead found for *dN/dS* (0.072) and for *dS* (0.481). This means that a statistically significant gradient in the rate of non-synonymous substitution is found along metabolic pathways with genes in top positions having experienced faster non-synonymous substitution rates.

Given that the top/bottom position implies directionality for the pathway, we repeated the analysis in the subset of 208 pathways that have no loops thus have a marked directional structure (Fig 3b and supplementary fig 4b). As before, KW tests show that there are significant differences between *dN/dS* (P=0.023) and *dN* (P=0.005) between the four classes considered, while no significant differences are found in *dS* (P=0.300). When looking at pairwise comparisons we find that *dN/dS* of genes of the one-step class have a significantly higher *dN/dS* than those belonging to the intermediate (P=0.035) and to the bottom (P=0.019) class, and the *dN* of genes belonging to the one-step class are significantly different from *dN* of the remaining three classes: top (P=0.034), intermediate (P=0.006), bottom (P=0.004).

In summary, the stringent KW test clearly highlighted differences between the rates of functional evolution of the genes belonging to the one-step class and those belonging to the other classes, pointing to their relaxed evolutionary constraint. This implies a gradient of non-synonymous substitution rates along metabolic pathways with genes at top-positions allowed to evolve at a faster rate than genes at intermediate and bottom positions, and with genes at intermediate and bottom positions being more tightly constrained to fix non-synonymous substitutions at a slower rate.

### Topological Measures of Positive Selected Genes

We measured the differences in the mean of the four centrality measures (in-degree, out-degree, closeness and betweenness) between genes with signatures of positive selection and the whole set of metabolic genes. We tested two sets: genes under positive selection with and without multiple testing correction. When only the six genes under positive selection after multiple testing correction were considered, no statistical significance is observed, even though genes under positive selection show higher out-degree and higher closeness than the average (0.33 vs. 0.22 for the out-degree and 0.38 vs. 0.28 for the closeness). Given the small number of positively selected genes (six genes), this same analysis was also repeated with all genes (49 genes) that have a p-value smaller than 0.05 in the positive selection test before the multiple test correction. The aim of this analysis is to test if there is any difference in the centrality measures values for the genes that belong to the tail of the positive selection test distribution. When these two groups are compared, higher out-degree (one tail permutation test, P = 0.0084) and higher closeness (one tail permutation test, P = 0.0116,) values are found for positively selected genes with an increase of 32% in the average value of out-degree and of 21% in closeness in genes under positive selection compared to the whole set of metabolic genes. This result shows that the tail of positive selection distribution is enriched by genes with higher out degree and higher closeness.

These results indicate that genes encoding enzymes with a greater number of reactions that make use of their products in the human metabolic pathways are more likely to present signals of positive selection than those with fewer enzymes using their products. Genes under positive selection have also higher closeness than the average and hence shorter path lengths to other nodes in the pathway.

## DISCUSSION

Linking the action of natural selection in the evolution of genes to the network structure and topology is an interesting approach to understand the constraints that the network structure may have on the evolution of complex molecular systems such as metabolism. Here, we have carried out a comprehensive molecular evolutionary study of human metabolism by investigating the selective pressures that acted on the enzyme-coding genes during the divergence of primates and rodents, and their relationships with functional and topological features of the pathways that constitute the system. Extensive studies have made possible the reconstruction of the biochemical pathways that constitute the metabolism; here we use this information to investigate the influence of the local network topology of the metabolic pathways in its evolutionary behavior by analyzing the distribution on the network of selective forces, be they in form of innovations (positive or adaptive selection) or in the strength of conservation (purifying selection).

The analysis of individual metabolic pathways instead of the whole metabolic network has several advantages: i) it allows the study of the influence onto the evolution of metabolic genes of their local relevant environment, that is, the context of the gene products in which the metabolic task is achieved; ii) it allows to separately study the functional units responsible for the different metabolic tasks, classify and compare them, and study the distribution of selective pressures within each functional unit; these functional units are particularly relevant because are likely to approximate the “molecular phenotypes” targeted by selection; and iii) it allows an intermediate analysis between the single enzyme and the whole metabolic network in the line of considering the hierarchical and topological structure of the pathways. However, the partitioning of the metabolic network into pathways is a somehow arbitrary process, the most arbitrary decision being the definition of pathway boundaries. Criteria for the definition of a pathway have been analyzed and compared (Caspi et al 2013; Green and Karp 2006) and the best collection of pathways available for human metabolism is comprised in the HumanCyc database (Romero et al. 2004; Caspi et al 2014). In HumanCyc, the criteria used for pathway definition are clearly stated and uniformly applied to the whole database (Caspi et al 2013). Importantly, the HumanCyc/MetaCyc approach of defining pathway boundaries through a multi-organism meta-metabolism implicitly introduces phylogenetic information and thus, pathways defined therein represent the best approximation of the functional units targeted by selection.

The final dataset under study was composed of 927 genes, whose products are integrated in 310 pathways. The species that have been considered are human, chimpanzee, gorilla, orangutan, mouse and rat, and thus the analysis embraces the divergence time of both primates and rodents. The tools for detecting positive selection and for measuring the strength of purifying selection (see methods) are based on the amino acid impact of nucleotide changes.

The detection of genes that underwent positive selection, and that are thus at the base of innovative changes, have resulted in a very small number of genes, six out of the 927 genes. These genes are: CYP2E1, a member of the cytochrome 450 family; HDC, a histidine decarboxylase; CES1, a carboxylesterase; DPM2, a subunit of the dolichol-phosphate mannose synthase complex; SPAM1, a hyaluronidase; and AKR1C1 an aldo-keto reductase. Two of the six genes that show sequence signature of positive selection, CYP2E1 and CES1, encode for detoxification enzymes and contribute to the solubility of molecules that must be expelled from the cell as fast as possible. Therefore, response to xenobiotic molecules has likely been a target of adaptive selection during primate and rodent divergence.

The main selective force in metabolism is the maintenance of the system through purifying selection; we have calculated this strength for each gene and analyzed the context according to biochemical and network properties. We observed a steep gradient of selective constraints that goes from genes serving functions within the inner core of intermediary metabolism (the most constrained), through those of a second layer of intermediary metabolism, to the outer peripheral layer of metabolism (*i.e.* secondary metabolism), which shows the most relaxed selective constraint. We have shown that the stronger selective constraint has acted on the genes whose encoded enzymes have roles within the inner core of metabolism; pathways comprised in this inner layer are involved in the transformations of small precursor metabolites for cell maintenance. These pathways are the oldest, the more phylogenetically conserved (Peregrin-Alvarez et al. 2003) and are enriched in enzymes exhibiting more substrate specificity (Nam et al. 2012). Accordingly, we have found that enzymes participating in more peripheral routes are evolutionarily less constrained and more plastic, having experienced faster functional evolution. Generalist enzymes, able to cope with a vast diversity of possible small molecular structures, populate pathways of this outer layer. The strategy of adopting generalist enzymes at the outer interface of metabolism may be a more efficient strategy than to develop a specific enzyme for each type of possible metabolite that may be present in the environment. Indeed, a global analysis of kinetic parameters of several thousands of known enzymes showed that central metabolism enzymes perform better in terms of catalytic constants (*i.e*. higher *k*_*cat*_ and *k_cat_/K_M_*) than secondary metabolism enzymes (Bar-Even et al 2011). In line with the authors’ suggestion, we have been able to prove that there is a stronger selective pressure on central metabolic enzymes, probably due to the need of maintaining the catalytic parameters that allow higher fluxes in central pathways, in comparison to those operating at lower fluxes and less specificity in secondary metabolism (Bar-Even et al 2011).

Isoenzymes are of special interest because they provide material for metabolic evolutionary innovation through sub- or neo-functionalization (Innan and Kondrashov 2010). Here we have found no differences in evolutionary rates between isoenzymes and the rest of the enzymes; no differences in their connectivities were found in the global metabolic network of *E.coli* (Light et al. 2005). Thus, both the sequence and the topological properties of isoenzymes do not differ from those of the rest of the enzymes. Selective constraint acting with the same strength onto these two classes of genes suggests that what may seem like alternative enzymes for the same metabolic function, are, indeed, equally “essential”, and the functional degeneracy is only apparent. Isoenzymes are likely to be essential to their function either through regulation, differential expression in time (different developmental stages), or in space, displaying their function in different cellular compartments or tissues. The study of evolutionary pressures over gene sequences has clearly pointed to lack of functional degeneracy for isoenzymes, given that redundancy leaves clear detectable footprints in terms of acceleration of substitution rates, as for example in the case of paralogous gene copies just after the duplication event (Innan and Kondrashov 2010).

To assess the influence of system properties on gene evolution we have used the local topology of the metabolic network. Among the many possible ways of representing metabolic pathways through graph structures, we have encoded pathways as reaction graphs, a type of representation in which nodes represent reactions; hence they can be directly associated to the genes that catalyze the reactions. In this representation edges represent metabolites (reaction substrate and products) and here we have considered their direction. So in our representation edges are indeed arrows and the resulting graph is a directed graph. By encoding metabolic pathways through directed graphs, we are able to take into account the physiological direction of the reactions in the cell.

Even in absence of specific information about the metabolic pathways for all the considered species, the relatively shallow phylogenetic divergence considered and the well-known conservation of metabolism among mammals ensures the use of a unique structure of pathways and function of all enzymes. The influence of the local network topology over gene’s evolution was investigated for both positive and purifying selection.

The influence of the local network topology over gene’s evolution had been previously investigated in few cases of specific metabolic pathways. The comprehensive analysis carried out here allowed revealing that this influence is pervasive and general patterns can be found. When positive selection has been considered we have found that positively selected genes have higher out-degree centralities than non-adaptive genes. Genes with high out-degree are not involved in the production of the final products of the metabolic task and, at the same time, are those whose products are subsequently transformed by a high fraction of different reactions in the pathway. Here we see that genes in these positions are preferentially targeted by adaptive evolution.

When purifying selection has been considered, in-degree centrality has been identified as the strongest topological factor constraining metabolic gene’s sequence evolution. Genes characterized by higher in-degree connectivities have evolved under stronger purifying selection. Within the context of biochemical reaction graphs in which each node represents a biochemical reaction, in-degree reflects the number of reactions that directly precede the given one, that is, the number of reactions that produce metabolites that are then taken as substrates by the given reaction. This means that in-degree centrality is higher for reaction with many incoming links and lower for reactions with few incoming links.

Besides the encoding of topological information through centrality measures we also used topological features that encode the position along the pathway (top, intermediate and bottom) of each enzyme. When this topological information was considered for plant biosynthetic pathways (Rausher et al. 1999; Lu and Rausher 2003; Rausher et al. 2008; Livingstone and Anderson 2009; Ramsay et al. 2009), a progressive relaxation of selective constraint along metabolic pathways was found. Here we found an opposite gradient when the whole set of human metabolic pathways were analyzed during the divergence of primates and rodents, with genes at top positions being under more relaxed constraint and genes at bottom position being under stronger selective constraint. Thus relaxed evolutionary forces at top positions allow broader evolutionary changes while, at the bottom, strong selective constraint narrows the allowed changes in functional variation, according to a funnel like model. This funnel like distribution of selective pressures along positions in the pathway is a general pattern found throughout the considered metabolic pathways. This result is consistent with the results obtained in the N-glycosylation pathway (Montanucci et al 2011) where genes at downstream positions of the pathways were found to be under stronger selective constraint than genes located at upstream (top) positions in the pathway which in turn had undergone more relaxed evolution. For metabolic routes that are directly connected with external environment, enzymes at top positions are usually more generalists in response to the need of using a broad diversity of possible metabolic material, while those at the bottom positions are more specialized, in response to the need of producing very specific products. It can be speculated that the results obtained here may reflect the relevance of accuracy in the synthesis of final metabolic products.

The comprehensive analysis of the whole set of human metabolic pathways revealed that both adaptive and purifying selection are not evenly distributed among the genes encoding the enzymes involved in metabolic pathways. Adaptive selection has targeted a small number of genes during the divergence of primates and rodents and adaptive genes are mainly involved in detoxification functions. Purifying selection has been a pervasive selective force dominating the evolution of metabolic genes during the divergence of primates and rodents. It has acted with different strengths according to the layer of metabolism over which it acts, with the inner core of metabolism being strongly conserved and with little or no room left for evolutionary innovation. From these results, it is tempting to conclude that it is less likely to innovate on pathways that were established in the early (*i.e*. prokaryotic) stages of evolution and that are involved in the synthesis of a small set of metabolic precursors linking the synthesis and degradation of essential biomolecules, namely sugars, lipids, amino acids and nucleotides. A more relaxed selection has been found for enzymes that manage higher levels of chemodiversity. This is the case of detoxification of xenobiotics or the biosynthesis of a wide spectrum of secondary metabolites that, by definition, are not directly involved in the survival of the organism, but rather in its ecological and behavioral traits.

## ACKNOWLEDGEMENTS

Authors would like to acknowledge Dr Peter Karp and Dr Ron Caspi for their support and help with data in HumanCyc database.

## Compliance with Ethical Standards

### Conflict of Interest

The authors declare that there is no conflict of interest

Research involving Human Participants and/or Animals: The research does not involve human or animal participants. It is solely based on databases.

### Informed consent: no need of informed consent

**Figure S1:**
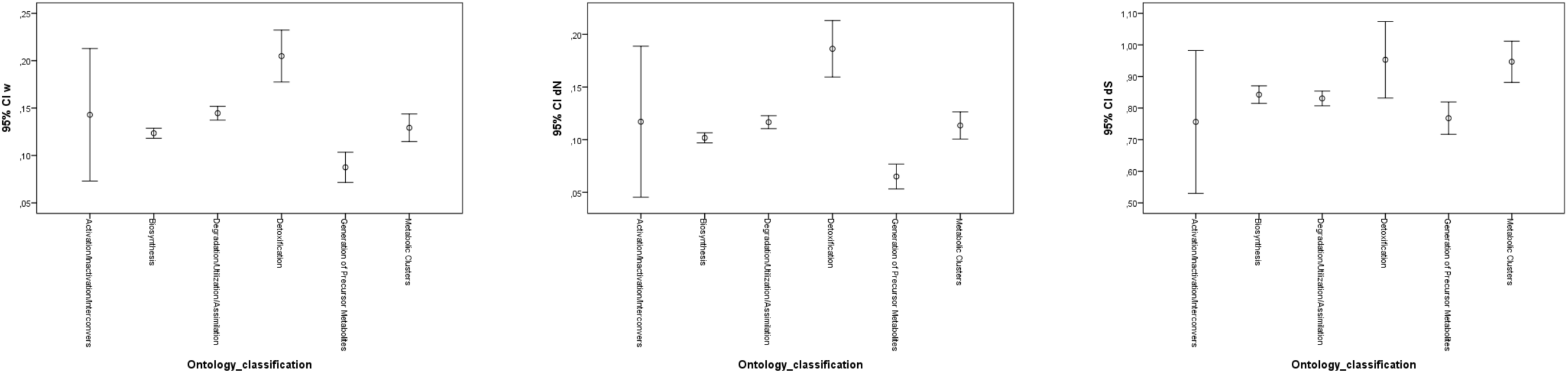
Graph representing *dN/dS*, *dN* and *dS* among genes belonging to different functional classes according to ontology-based classification.

**Figure S2:**
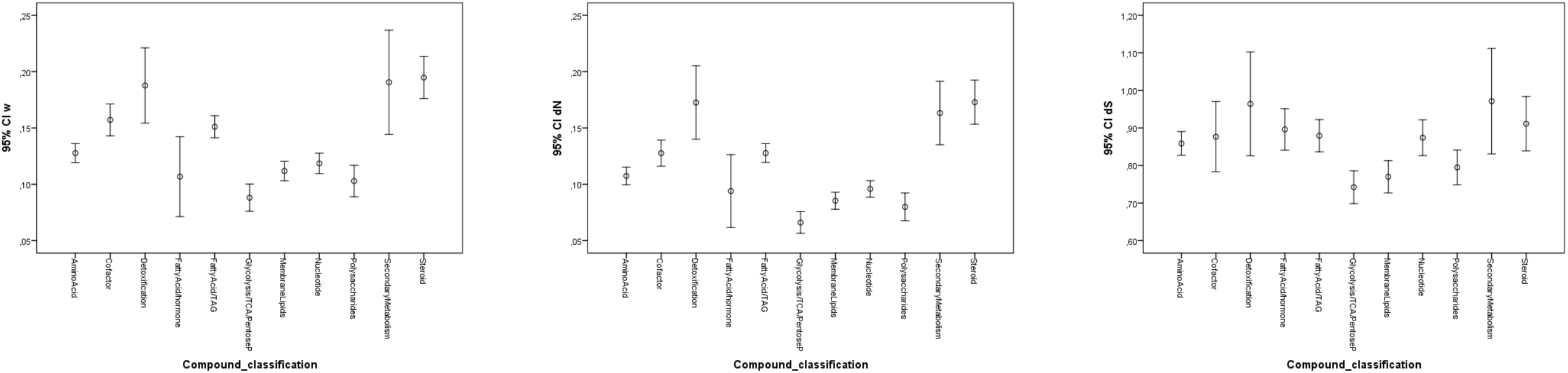
Graph representing *dN/dS*, *dN* and *dS* among genes belonging to different functional classes according to compound-based classification.

**Figure S3:**
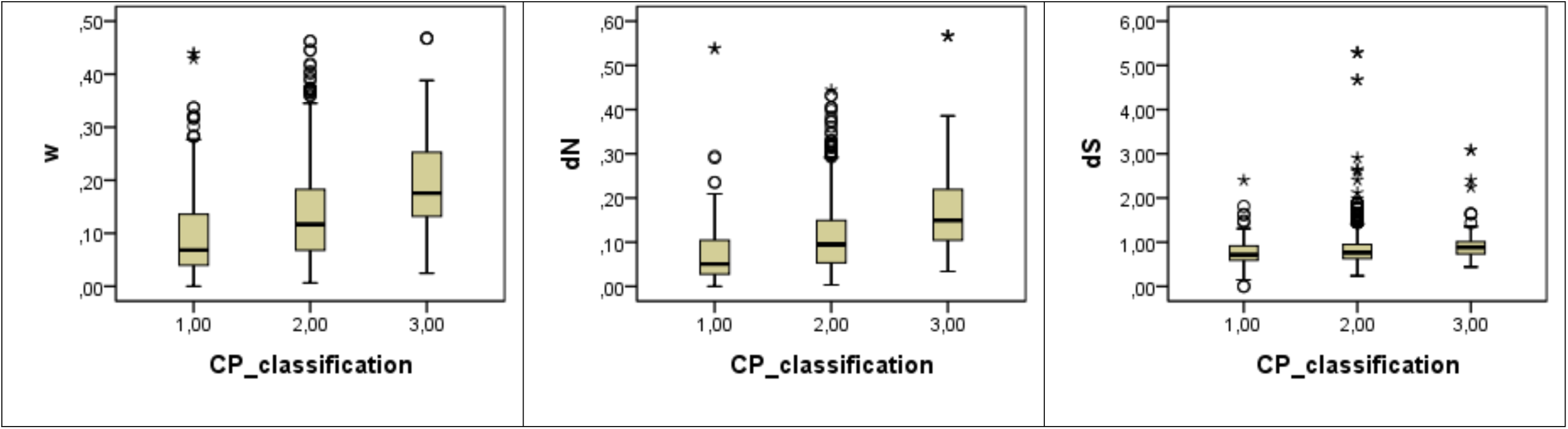
Boxplots representing ω (*dN/dS*), *dN* and *dS* among genes belonging to different functional classes. Class 1 comprises the inner metabolism (Glycosis/TCA/PentoseP, Polysaccharides), class 2 comprises the second layer (Membrane Lipids metabolism, nucleotide metabolism, Fatty acid/TAG, Cofactor, Fatty Acid/hormone, and Aminoacid) while class 3 comprises the outer layer of cell metabolism (Steroid, Secondary Metabolism and Detoxification).. Boxes are 25th and 75th quartiles, black bar within the box represents the median, whiskers indicate minimum and maximum and dots and stars represent most extreme data point higher than 1.5 interquartile range from box.

**Figure S4:**
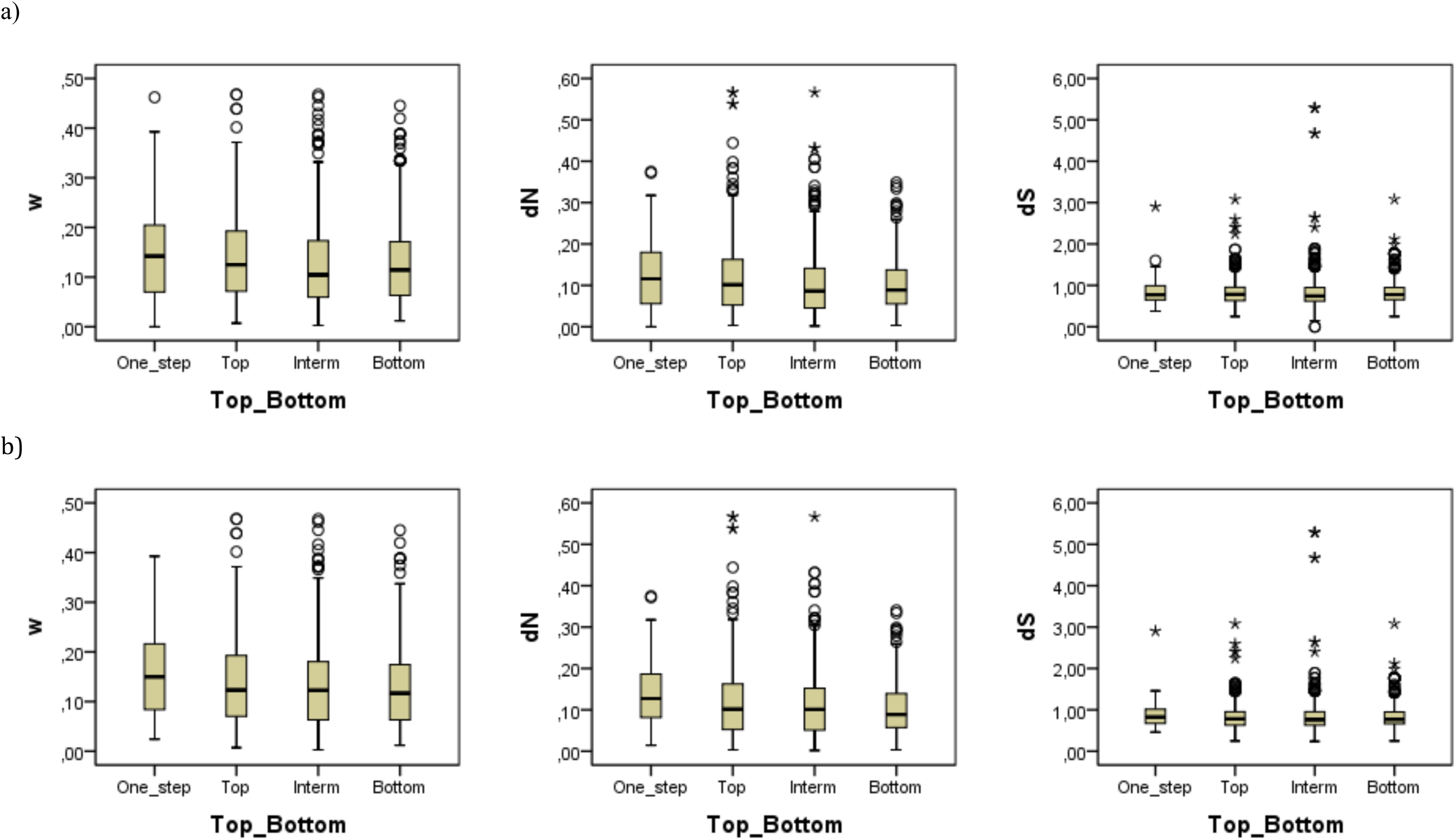
**a)** Boxplots representing *dN/dS*, *dN* and *dS* among genes whose position within the pathway belongs to four classes (one-step, top, intermediate, bottom positions) for the 275 base pathways. **b)** The same for the 208 base pathways with no loops. Dots show the mean ± 2 standard error (SE). Boxes are 25th and 75th quartiles, black bar within the box represents the median, whiskers indicate minimum and maximum and dots and stars represent most extreme data point higher than 1.5 interquartile range from the box.

**Table S1.** List of the 310 pathways, their codes, common names and their classifications.

| **PATHWAY CODE** | **PATHWAY COMMON NAME** | **ONTOLOGY-BASED CLASSIFICATION** | **COMPOUND-BASED CLASSIFICATION** |
| --- | --- | --- | --- |
| 2PHENDEG-PWY | phenylethylamine degradation I | Degradation/Utilization/Assimilation | AminoAcid |
| ADENOSYLHOMOCYSCAT-PWY | methionine salvage | Biosynthesis | AminoAcid |
| ALANINE-DEG3-PWY | alanine biosynthesis/degradation | Degradation/Utilization/Assimilation | AminoAcid |
| ARGININE-SYN4-PWY | ornithine *de novo* biosynthesis | Biosynthesis | AminoAcid |
| ARGSPECAT-PWY | spermine biosynthesis | Biosynthesis | AminoAcid |
| ASPARAGINE-BIOSYNTHESIS | asparagine biosynthesis | Biosynthesis | AminoAcid |
| ASPARAGINE-DEG1-PWY | asparagine degradation | Degradation/Utilization/Assimilation | AminoAcid |
| ASPARTATESYN-PWY | aspartate biosynthesis | Biosynthesis | AminoAcid |
| BETA-ALA-DEGRADATION-I-PWY | β-alanine degradation | Degradation/Utilization/Assimilation | AminoAcid |
| BSUBPOLYAMSYN-PWY | spermidine biosynthesis | Biosynthesis | AminoAcid |
| CYSTEINE-DEG-PWY | L-cysteine degradation I | Degradation/Utilization/Assimilation | AminoAcid |
| GLNSYN-PWY | glutamine biosynthesis | Biosynthesis | AminoAcid |
| GLUDEG-I-PWY | GABA shunt | Degradation/Utilization/Assimilation | AminoAcid |
| GLUTAMATE-SYN2-PWY | glutamate biosynthesis/degradation | Biosynthesis | AminoAcid |
| GLUTAMINDEG-PWY | glutamine degradation/glutamate biosynthesis | Degradation/Utilization/Assimilation | AminoAcid |
| GLUTATHIONESYN-PWY | glutathione biosynthesis | Biosynthesis | AminoAcid |
| GLYCLEAV-PWY | glycine cleavage | Degradation/Utilization/Assimilation | AminoAcid |
| GLYSYN-ALA-PWY | glycine biosynthesis | Biosynthesis | AminoAcid |
| GLYSYN-PWY | glycine/serine biosynthesis | Biosynthesis | AminoAcid |
| HOMOCYSDEGR-PWY | cysteine biosynthesis/homocysteine degradation (trans-sulfuration) | Degradation/Utilization/Assimilation | AminoAcid |
| HYDROXYPRODEG-PWY | 4-hydroxyproline degradation | Degradation/Utilization/Assimilation | AminoAcid |
| ILEUDEG-PWY | isoleucine degradation | Degradation/Utilization/Assimilation | AminoAcid |
| LEU-DEG2-PWY | leucine degradation | Degradation/Utilization/Assimilation | AminoAcid |
| LYSINE-DEG1-PWY | lysine degradation I (saccharopine pathway) | Degradation/Utilization/Assimilation | AminoAcid |
| METHIONINE-DEG1-PWY | methionine degradation | Degradation/Utilization/Assimilation | AminoAcid |
| PHENYLALANINE-DEG1-PWY | phenylalanine degradation/tyrosine biosynthesis | Degradation/Utilization/Assimilation | AminoAcid |
| PROSYN-PWY | proline biosynthesis | Biosynthesis | AminoAcid |
| PROUT-PWY | proline degradation | Degradation/Utilization/Assimilation | AminoAcid |
| PWY-0 | putrescine degradation III | Degradation/Utilization/Assimilation | AminoAcid |
| PWY0-1305 | glutamate dependent acid resistance | Detoxification | AminoAcid |
| PWY-3661-1 | glycine betaine degradation | Degradation/Utilization/Assimilation | AminoAcid |
| PWY-40 | putrescine biosynthesis II | Biosynthesis | AminoAcid |
| PWY-4041 | γ-glutamyl cycle | Superpathways | AminoAcid |
| PWY-4061 | glutathione-mediated detoxification | Detoxification | AminoAcid |
| PWY-4081 | glutathione redox reactions I | Biosynthesis | AminoAcid |
| PWY-46 | putrescine biosynthesis I | Biosynthesis | AminoAcid |
| PWY-4921 | protein citrullination | Biosynthesis | AminoAcid |
| PWY-4983 | citrulline-nitric oxide cycle | Degradation/Utilization/Assimilation | AminoAcid |
| PWY-4984 | urea cycle | Degradation/Utilization/Assimilation | AminoAcid |
| PWY-5030 | histidine degradation | Degradation/Utilization/Assimilation | AminoAcid |
| PWY-5046 | 2-oxoisovalerate decarboxylation to isobutanoyl-CoA | Generation of Precursor Metabolites and Energy | AminoAcid |
| PWY-5177 | glutaryl-CoA degradation | Degradation/Utilization/Assimilation | AminoAcid |
| PWY-5326 | sulfite oxidation | Degradation/Utilization/Assimilation | AminoAcid |
| PWY-5328 | superpathway of methionine degradation | Degradation/Utilization/Assimilation | AminoAcid |
| PWY-5329 | L-cysteine degradation II | Degradation/Utilization/Assimilation | AminoAcid |
| PWY-5331 | taurine biosynthesis | Biosynthesis | AminoAcid |
| PWY-5340 | sulfate activation for sulfonation | Degradation/Utilization/Assimilation | AminoAcid |
| PWY-5350 | thiosulfate disproportionation III (rhodanese) | Degradation/Utilization/Assimilation | AminoAcid |
| PWY-5651 | tryptophan degradation to 2-amino-3-carboxymuconate semialdehyde | Degradation/Utilization/Assimilation | AminoAcid |
| PWY-5652 | 2-amino-3-carboxymuconate semialdehyde degradation to glutaryl-CoA | Degradation/Utilization/Assimilation | AminoAcid |
| PWY-5905 | hypusine biosynthesis | Biosynthesis | AminoAcid |
| PWY-5921 | L-glutamine tRNA biosynthesis | Biosynthesis | AminoAcid |
| PWY-6030 | serotonin and melatonin biosynthesis | Biosynthesis | AminoAcid |
| PWY-6100 | L-carnitine biosynthesis | Biosynthesis | AminoAcid |
| PWY-6117 | spermine and spermidine degradation I | Degradation/Utilization/Assimilation | AminoAcid |
| PWY-6133 | (S)-reticuline biosynthesis | Biosynthesis | AminoAcid |
| PWY-6158 | creatine-phosphate biosynthesis | Biosynthesis | AminoAcid |
| PWY-6173 | histamine biosynthesis | Biosynthesis | AminoAcid |
| PWY-6181 | histamine degradation | Degradation/Utilization/Assimilation | AminoAcid |
| PWY-6241 | thyroid hormone biosynthesis | Biosynthesis | AminoAcid |
| PWY-6260 | thyroid hormone metabolism I (via deiodination) | Degradation/Utilization/Assimilation | AminoAcid |
| PWY-6261 | thyroid hormone metabolism II (via conjugation and/or degradation) | Degradation/Utilization/Assimilation | AminoAcid |
| PWY-6281 | selenocysteine biosynthesis | Biosynthesis | AminoAcid |
| PWY-6307 | tryptophan degradation X (mammalian, via tryptamine) | Degradation/Utilization/Assimilation | AminoAcid |
| PWY-6313 | serotonin degradation | Degradation/Utilization/Assimilation | AminoAcid |
| PWY-6334 | L-dopa degradation | Degradation/Utilization/Assimilation | AminoAcid |
| PWY-6342 | noradrenaline and adrenaline degradation | Degradation/Utilization/Assimilation | AminoAcid |
| PWY-6481 | L-dopachrome biosynthesis | Biosynthesis | AminoAcid |
| PWY-6482 | diphthamide biosynthesis | Biosynthesis | AminoAcid |
| PWY-6498 | eumelanin biosynthesis | Biosynthesis | AminoAcid |
| PWY66-301 | catecholamine biosynthesis | Biosynthesis | AminoAcid |
| PWY66-425 | lysine degradation II (pipecolate pathway) | Degradation/Utilization/Assimilation | AminoAcid |
| PWY66-426 | hydrogen sulfide biosynthesis (trans-sulfuration) | Metabolic Clusters | AminoAcid |
| PWY66-428 | threonine degradation | Degradation/Utilization/Assimilation | AminoAcid |
| PWY6666-2 | dopamine degradation | Degradation/Utilization/Assimilation | AminoAcid |
| PWY-6688 | thyronamine and iodothyronamine metabolism | Degradation/Utilization/Assimilation | AminoAcid |
| PWY-6755 | *S*-methyl-5-thio-α-D-ribose 1-phosphate degradation | Degradation/Utilization/Assimilation | AminoAcid |
| PWY-6756 | *S*-methyl-5'-thioadenosine degradation | Degradation/Utilization/Assimilation | AminoAcid |
| SAM-PWY | S-adenosyl-L-methionine biosynthesis | Biosynthesis | AminoAcid |
| SERDEG-PWY | L-serine degradation | Degradation/Utilization/Assimilation | AminoAcid |
| SER-GLYSYN-PWY-1 | serine and glycine biosynthesis | Superpathways | AminoAcid |
| SERSYN-PWY | serine biosynthesis (phosphorylated route) | Biosynthesis | AminoAcid |
| TRYPTOPHAN-DEGRADATION-1 | tryptophan degradation | Degradation/Utilization/Assimilation | AminoAcid |
| TYRFUMCAT-PWY | tyrosine degradation | Degradation/Utilization/Assimilation | AminoAcid |
| VALDEG-PWY | valine degradation | Degradation/Utilization/Assimilation | AminoAcid |
| PWY66-414 | superpathway of choline degradation to L-serine | Superpathways | AminoAcid |
| PWY-6292 | cysteine biosynthesis | Superpathways | AminoAcid |
| PWY66-401 | superpathway of tryptophan utilization | Superpathways | AminoAcid |
| COA-PWY-1 | coenzyme A biosynthesis | Biosynthesis | Cofactor |
| GLYCGREAT-PWY | creatine biosynthesis | Degradation/Utilization/Assimilation | Cofactor |
| HEME-BIOSYNTHESIS-II | heme biosynthesis from uroporphyrinogen-III I | Biosynthesis | Cofactor |
| NAD-BIOSYNTHESIS-III | NAD salvage | Biosynthesis | Cofactor |
| NADPHOS-DEPHOS-PWY-1 | NAD phosphorylation and dephosphorylation | Biosynthesis | Cofactor |
| NADSYN-PWY | NAD *de novo* biosynthesis | Biosynthesis | Cofactor |
| PLPSAL-PWY | pyridoxal 5'-phosphate salvage | Biosynthesis | Cofactor |
| PWY0-1264 | biotin-carboxyl carrier protein assembly | Biosynthesis | Cofactor |
| PWY0-1275 | lipoate biosynthesis and incorporation | Biosynthesis | Cofactor |
| PWY0-522 | lipoate salvage | Biosynthesis | Cofactor |
| PWY-2161 | folate polyglutamylaton | Biosynthesis | Cofactor |
| PWY-2161B | glutamate removal from folates | Biosynthesis | Cofactor |
| PWY-2201 | folate transformations | Biosynthesis | Cofactor |
| PWY-5189 | tetrapyrrole biosynthesis | Biosynthesis | Cofactor |
| PWY-5653 | NAD biosynthesis from 2-amino-3-carboxymuconate semialdehyde | Biosynthesis | Cofactor |
| PWY-5663 | tetrahydrobiopterin *de novo* biosynthesis | Biosynthesis | Cofactor |
| PWY-5754 | 4-hydroxybenzoate biosynthesis | Biosynthesis | Cofactor |
| PWY-5872 | ubiquinol-10 biosynthesis | Biosynthesis | Cofactor |
| PWY-5874 | heme degradation | Degradation/Utilization/Assimilation | Cofactor |
| PWY-5963 | thio-molybdenum cofactor biosynthesis | Biosynthesis | Cofactor |
| PWY-6076 | 1,25-dihydroxyvitamin D<sub>3</sub> biosynthesis | Biosynthesis | Cofactor |
| PWY-6309 | L-kynurenine degradation | Superpathways | Cofactor |
| PWY-6430 | thymine degradation | Degradation/Utilization/Assimilation | Cofactor |
| PWY-6613 | tetrahydrofolate salvage from 5,10-methenyltetrahydrofolate | Biosynthesis | Cofactor |
| PWY66-201 | nicotine degradation IV | Degradation/Utilization/Assimilation | Cofactor |
| PWY66-221 | nicotine degradation III | Degradation/Utilization/Assimilation | Cofactor |
| PWY66-366 | flavin biosynthesis | Biosynthesis | Cofactor |
| PWY-6823 | molybdenum cofactor biosynthesis | Biosynthesis | Cofactor |
| PWY-6857 | retinol biosynthesis | Biosynthesis | Cofactor |
| PWY-6872 | retinoate biosynthesis I | Biosynthesis | Cofactor |
| PWY-6875 | retinoate biosynthesis II | Biosynthesis | Cofactor |
| PWY-6898 | thiamin salvage III | Biosynthesis | Cofactor |
| PWY-6938 | NADH repair | Generation of Precursor Metabolites and Energy | Cofactor |
| PWY-7250 | iron-sulfur cluster biosynthesis | Biosynthesis | Cofactor |
| PWY-7283 | wybutosine biosynthesis | Superpathways | Cofactor |
| PWY-7286 | 7-(3-amino-3-carboxypropyl)-wyosine biosynthesis | Biosynthesis | Cofactor |
| THIOREDOX-PWY | thioredoxin pathway | Biosynthesis | Cofactor |
| PWY-5920 | heme biosynthesis | Superpathways | Cofactor |
| DETOX1-PWY | superoxide radicals degradation | Detoxification | Detoxification |
| GLUT-REDOX-PWY | glutathione redox reactions II | Biosynthesis | Detoxification |
| MGLDLCTANA-PWY | methylglyoxal degradation VI | Detoxification | Detoxification |
| PWY-1801 | formaldehyde oxidation | Degradation/Utilization/Assimilation | Detoxification |
| PWY-4202 | arsenate detoxification I (glutaredoxin) | Detoxification | Detoxification |
| PWY-5386 | methylglyoxal degradation I | Detoxification | Detoxification |
| PWY-5453 | methylglyoxal degradation III | Detoxification | Detoxification |
| PWY-6502 | oxidized GTP and dGTP detoxification | Metabolic Clusters | Detoxification |
| PWY66-241 | bupropion degradation | Degradation/Utilization/Assimilation | Detoxification |
| PWY-7112 | 4-hydroxy-2-nonenal detoxification | Detoxification | Detoxification |
| PWY66-392 | lipoxin biosynthesis | Biosynthesis | FattyAcid/hormone |
| PWY66-393 | aspirin-triggered lipoxin biosynthesis | Biosynthesis | FattyAcid/hormone |
| PWY66-394 | aspirin triggered resolvin E biosynthesis | Biosynthesis | FattyAcid/hormone |
| PWY66-395 | aspirin triggered resolvin D biosynthesis | Biosynthesis | FattyAcid/hormone |
| PWY66-397 | resolvin D biosynthesis | Biosynthesis | FattyAcid/hormone |
| FAO-PWY | fatty acid β-oxidation | Degradation/Utilization/Assimilation | FattyAcid/TAG |
| FASYN-ELONG-PWY | fatty acid elongation -- saturated | Biosynthesis | FattyAcid/TAG |
| LIPAS-PWY | triacylglycerol degradation | Degradation/Utilization/Assimilation | FattyAcid/TAG |
| LIPASYN-PWY | phospholipases | Metabolic Clusters | FattyAcid/TAG |
| PROPIONMET-PWY | propionyl-CoA degradation | Degradation/Utilization/Assimilation | FattyAcid/TAG |
| PWY-5130 | 2-oxobutanoate degradation | Superpathways | FattyAcid/TAG |
| PWY-5137 | fatty acid β-oxidation (unsaturated, odd number) | Degradation/Utilization/Assimilation | FattyAcid/TAG |
| PWY-5143 | fatty acid activation | Activation/Inactivation/Interconversion | FattyAcid/TAG |
| PWY-5148 | acyl-CoA hydrolysis | Biosynthesis | FattyAcid/TAG |
| PWY-5451 | acetone degradation I (to methylglyoxal) | Degradation/Utilization/Assimilation | FattyAcid/TAG |
| PWY-5966-1 | fatty acid biosynthesis initiation | Biosynthesis | FattyAcid/TAG |
| PWY-5972 | stearate biosynthesis | Biosynthesis | FattyAcid/TAG |
| PWY-5994 | palmitate biosynthesis | Biosynthesis | FattyAcid/TAG |
| PWY-5996 | oleate biosynthesis | Biosynthesis | FattyAcid/TAG |
| PWY-6000 | γ-linolenate biosynthesis | Biosynthesis | FattyAcid/TAG |
| PWY-6012-1 | acyl carrier protein metabolism | Biosynthesis | FattyAcid/TAG |
| PWY-6111 | mitochondrial L-carnitine shuttle | Degradation/Utilization/Assimilation | FattyAcid/TAG |
| PWY-6535 | 4-aminobutyrate degradation | Degradation/Utilization/Assimilation | FattyAcid/TAG |
| PWY66-161 | oxidative ethanol degradation III | Degradation/Utilization/Assimilation | FattyAcid/TAG |
| PWY66-162 | ethanol degradation IV | Degradation/Utilization/Assimilation | FattyAcid/TAG |
| PWY66-21 | ethanol degradation II | Degradation/Utilization/Assimilation | FattyAcid/TAG |
| PWY66-374 | C20 prostanoid biosynthesis | Biosynthesis | FattyAcid/TAG |
| PWY66-375 | leukotriene biosynthesis | Biosynthesis | FattyAcid/TAG |
| PWY66-387 | fatty acid α-oxidation | Degradation/Utilization/Assimilation | FattyAcid/TAG |
| PWY66-388 | fatty acid α-oxidation III | Degradation/Utilization/Assimilation | FattyAcid/TAG |
| PWY66-389 | phytol degradation | Degradation/Utilization/Assimilation | FattyAcid/TAG |
| PWY66-391 | fatty acid β-oxidation (peroxisome) | Degradation/Utilization/Assimilation | FattyAcid/TAG |
| PWY6666-1 | anandamide degradation | Degradation/Utilization/Assimilation | FattyAcid/TAG |
| PWY-7049 | eicosapentaenoate biosynthesis | Biosynthesis | FattyAcid/TAG |
| TRIGLSYN-PWY | triacylglycerol biosynthesis | Biosynthesis | FattyAcid/TAG |
| MALATE-ASPARTATE-SHUTTLE-PWY | malate-aspartate shuttle | Degradation/Utilization/Assimilation | Glycolysis/TCA/PentoseP |
| NONOXIPENT-PWY | pentose phosphate pathway (non-oxidative branch) | Generation of Precursor Metabolites and Energy | Glycolysis/TCA/PentoseP |
| OXIDATIVEPENT-PWY-1 | pentose phosphate pathway (oxidative branch) | Generation of Precursor Metabolites and Energy | Glycolysis/TCA/PentoseP |
| PWY0-1313 | acetate conversion to acetyl-CoA | Degradation/Utilization/Assimilation | Glycolysis/TCA/PentoseP |
| PWY0-662 | PRPP biosynthesis | Biosynthesis | Glycolysis/TCA/PentoseP |
| PWY-4261 | glycerol degradation | Degradation/Utilization/Assimilation | Glycolysis/TCA/PentoseP |
| PWY-5084 | 2-oxoglutarate decarboxylation to succinyl-CoA | Degradation/Utilization/Assimilation | Glycolysis/TCA/PentoseP |
| PWY-5172 | acetyl-CoA biosynthesis from citrate | Generation of Precursor Metabolites and Energy | Glycolysis/TCA/PentoseP |
| PWY-5481 | lactate fermentation (reoxidation of cytosolic NADH) | Generation of Precursor Metabolites and Energy | Glycolysis/TCA/PentoseP |
| PWY-6118 | glycerol-3-phosphate shuttle | Generation of Precursor Metabolites and Energy | Glycolysis/TCA/PentoseP |
| PWY-6405 | Rapoport-Luebering glycolytic shunt | Biosynthesis | Glycolysis/TCA/PentoseP |
| PWY66-367 | ketogenesis | Generation of Precursor Metabolites and Energy | Glycolysis/TCA/PentoseP |
| PWY66-368 | ketolysis | Generation of Precursor Metabolites and Energy | Glycolysis/TCA/PentoseP |
| PWY66-398 | TCA cycle | Generation of Precursor Metabolites and Energy | Glycolysis/TCA/PentoseP |
| PWY66-399 | gluconeogenesis | Biosynthesis | Glycolysis/TCA/PentoseP |
| PWY66-400 | glycolysis | Generation of Precursor Metabolites and Energy | Glycolysis/TCA/PentoseP |
| PWY66-423 | fructose 2,6-bisphosphate synthesis/dephosphorylation | Biosynthesis | Glycolysis/TCA/PentoseP |
| PYRUVDEHYD-PWY | pyruvate decarboxylation to acetyl CoA | Generation of Precursor Metabolites and Energy | Glycolysis/TCA/PentoseP |
| PENTOSE-P-PWY | pentose phosphate pathway | Superpathways | Glycolysis/TCA/PentoseP |
| PWY66-407 | superpathway of conversion of glucose to acetyl CoA and entry into the TCA cycle | Superpathways | Glycolysis/TCA/PentoseP |
| CHOLINE-BETAINE-ANA-PWY | choline degradation | Degradation/Utilization/Assimilation | MembraneLipids |
| MANNOSYL-CHITO-DOLICHOL-BIOSYNTHESIS | dolichyl-diphosphooligosaccharide biosynthesis | Biosynthesis | MembraneLipids |
| PWY-2301 | *myo*-inositol *de novo* biosynthesis | Biosynthesis | MembraneLipids |
| PWY3DJ-11281 | sphingomyelin metabolism/ceramide salvage | Biosynthesis | MembraneLipids |
| PWY3DJ-11470 | sphingosine and sphingosine-1-phosphate metabolism | Degradation/Utilization/Assimilation | MembraneLipids |
| PWY3DJ-12 | ceramide *de novo* biosynthesis | Biosynthesis | MembraneLipids |
| PWY3O-450 | phosphatidylcholine biosynthesis | Biosynthesis | MembraneLipids |
| PWY4FS-6 | phosphatidylethanolamine biosynthesis II | Biosynthesis | MembraneLipids |
| PWY-5269 | cardiolipin biosynthesis | Biosynthesis | MembraneLipids |
| PWY-5667 | CDP-diacylglycerol biosynthesis | Biosynthesis | MembraneLipids |
| PWY-6129 | dolichol and dolichyl phosphate biosynthesis | Biosynthesis | MembraneLipids |
| PWY-6351 | D-*myo*-inositol (1,4,5)-trisphosphate biosynthesis | Biosynthesis | MembraneLipids |
| PWY-6352 | 3-phosphoinositide biosynthesis | Biosynthesis | MembraneLipids |
| PWY-6362 | 1D-*myo*-inositol hexakisphosphate biosynthesis II (mammalian) | Biosynthesis | MembraneLipids |
| PWY-6363 | D-*myo*-inositol (1,4,5)-trisphosphate degradation | Biosynthesis | MembraneLipids |
| PWY-6364 | D-*myo*-inositol (1,3,4)-trisphosphate biosynthesis | Biosynthesis | MembraneLipids |
| PWY-6365 | D-*myo*-inositol (3,4,5,6)-tetrakisphosphate biosynthesis | Biosynthesis | MembraneLipids |
| PWY-6366 | D-*myo*-inositol (1,4,5,6)-tetrakisphosphate biosynthesis | Biosynthesis | MembraneLipids |
| PWY-6367 | D-*myo*-inositol-5-phosphate metabolism | Biosynthesis | MembraneLipids |
| PWY-6368 | 3-phosphoinositide degradation | Degradation/Utilization/Assimilation | MembraneLipids |
| PWY-6369 | inositol pyrophosphates biosynthesis | Biosynthesis | MembraneLipids |
| PWY-6554 | 1D-*myo*-inositol hexakisphosphate biosynthesis V (from Ins(1,3,4)P3) | Biosynthesis | MembraneLipids |
| PWY-7501 | phosphatidylserine biosynthesis I | Biosynthesis | MembraneLipids |
| PWY-6358 | superpathway of D-*myo*-inositol (1,4,5)-trisphosphate metabolism | Superpathways | MembraneLipids |
| PWY-6371 | superpathway of inositol phosphate compounds | Superpathways | MembraneLipids |
| P121-PWY | adenine and adenosine salvage I | Biosynthesis | Nucleotide |
| PWY0-1295 | pyrimidine ribonucleosides degradation | Degradation/Utilization/Assimilation | Nucleotide |
| PWY0-1296 | purine ribonucleosides degradation to ribose-1-phosphate | Degradation/Utilization/Assimilation | Nucleotide |
| PWY-3982 | uracil degradation | Biosynthesis | Nucleotide |
| PWY-5686 | UMP biosynthesis | Biosynthesis | Nucleotide |
| PWY-5695 | urate biosynthesis/inosine 5'-phosphate degradation | Degradation/Utilization/Assimilation | Nucleotide |
| PWY-6121 | 5-aminoimidazole ribonucleotide biosynthesis | Biosynthesis | Nucleotide |
| PWY-6124 | inosine-5'-phosphate biosynthesis | Biosynthesis | Nucleotide |
| PWY-6608 | guanosine nucleotides degradation | Degradation/Utilization/Assimilation | Nucleotide |
| PWY-6609 | adenine and adenosine salvage III | Biosynthesis | Nucleotide |
| PWY-6619 | adenine and adenosine salvage II | Biosynthesis | Nucleotide |
| PWY-6620 | guanine and guanosine salvage | Biosynthesis | Nucleotide |
| PWY66-385 | dTMP *<i>de novo</i>* biosynthesis (mitochondrial) | Biosynthesis | Nucleotide |
| PWY66-420 | carnosine biosynthesis | Biosynthesis | Nucleotide |
| PWY66-421 | homocarnosine biosynthesis | Biosynthesis | Nucleotide |
| PWY-7176 | UTP and CTP *<i>de novo</i>* biosynthesis | Biosynthesis | Nucleotide |
| PWY-7177 | UTP and CTP dephosphorylation II | Degradation/Utilization/Assimilation | Nucleotide |
| PWY-7179-1 | purine deoxyribonucleosides degradation | Degradation/Utilization/Assimilation | Nucleotide |
| PWY-7180 | 2'-deoxy-α-D-ribose 1-phosphate degradation | Degradation/Utilization/Assimilation | Nucleotide |
| PWY-7181 | pyrimidine deoxyribonucleosides degradation | Degradation/Utilization/Assimilation | Nucleotide |
| PWY-7184 | pyrimidine deoxyribonucleotides *<i>de novo</i>* biosynthesis | Metabolic Clusters | Nucleotide |
| PWY-7185 | UTP and CTP dephosphorylation I | Degradation/Utilization/Assimilation | Nucleotide |
| PWY-7193 | pyrimidine ribonucleosides salvage I | Biosynthesis | Nucleotide |
| PWY-7197 | pyrimidine deoxyribonucleotide phosphorylation | Metabolic Clusters | Nucleotide |
| PWY-7199 | pyrimidine deoxyribonucleosides salvage | Biosynthesis | Nucleotide |
| PWY-7205 | CMP phosphorylation | Biosynthesis | Nucleotide |
| PWY-7210 | pyrimidine deoxyribonucleotides biosynthesis from CTP | Metabolic Clusters | Nucleotide |
| PWY-7219 | adenosine ribonucleotides *<i>de novo</i>* biosynthesis | Biosynthesis | Nucleotide |
| PWY-7221 | guanosine ribonucleotides *<i>de novo</i>* biosynthesis | Biosynthesis | Nucleotide |
| PWY-7224 | purine deoxyribonucleosides salvage | Metabolic Clusters | Nucleotide |
| PWY-7226 | guanosine deoxyribonucleotides *<i>de novo</i>* biosynthesis | Biosynthesis | Nucleotide |
| PWY-7227 | adenosine deoxyribonucleotides *<i>de novo</i>* biosynthesis | Biosynthesis | Nucleotide |
| SALVADEHYPOX-PWY | adenosine nucleotides degradation | Degradation/Utilization/Assimilation | Nucleotide |
| PWY0-162 | superpathway of pyrimidine ribonucleotides *<i>de novo</i>* biosynthesis | Superpathways | Nucleotide |
| PWY-841 | purine nucleotides *<i>de novo</i>* biosynthesis | Superpathways | Nucleotide |
| PWY-6353 | purine nucleotides degradation | Superpathways | Nucleotide |
| PWY-7209 | pyrimidine ribonucleosides degradation | Superpathways | Nucleotide |
| PWY-7200 | superpathway of pyrimidine deoxyribonucleoside salvage | Superpathways | Nucleotide |
| PWY-7228 | guanosine nucleotides *<i>de novo</i>* biosynthesis | Superpathways | Nucleotide |
| PWY-7211 | superpathway of pyrimidine deoxyribonucleotides<br>*de novo* biosynthesis | Superpathways | Nucleotide |
| PWY66-409 | superpathway of purine nucleotide salvage | Superpathways | Nucleotide |
| BGALACT-PWY | lactose degradation III | Degradation/Utilization/Assimilation | Polysaccharides |
| GLUAMCAT-PWY | *N*-acetylglucosamine degradation I | Degradation/Utilization/Assimilation | Polysaccharides |
| MANNCAT-PWY | D-mannose degradation | Degradation/Utilization/Assimilation | Polysaccharides |
| PWY0-1182 | trehalose degradation | Degradation/Utilization/Assimilation | Polysaccharides |
| PWY-4101 | sorbitol degradation I | Degradation/Utilization/Assimilation | Polysaccharides |
| PWY-4821 | UDP-D-xylose and UDP-D-glucuronate<br>biosynthesis | Biosynthesis | Polysaccharides |
| PWY-5067 | glycogen biosynthesis | Biosynthesis | Polysaccharides |
| PWY-5512 | UDP-*N*-acetyl-D-galactosamine<br>biosynthesis I | Biosynthesis | Polysaccharides |
| PWY-5514 | UDP-*N*-acetyl-D-galactosamine<br>biosynthesis II | Biosynthesis | Polysaccharides |
| PWY-55y5 | D-glucuronate degradation | Degradation/Utilization/Assimilation | Polysaccharides |
| PWY-5659 | GDP-mannose biosynthesis | Biosynthesis | Polysaccharides |
| PWY-5661-1 | GDP-glucose biosynthesis II | Biosynthesis | Polysaccharides |
| PWY-5941-1 | glycogenolysis | Degradation/Utilization/Assimilation | Polysaccharides |
| PWY-6 | GDP-L-fucose biosynthesis II (from L-fucose) | Biosynthesis | Polysaccharides |
| PWY-6138 | CMP-*N*-acetylneuraminate biosynthesis I<br>(eukaryotes) | Biosynthesis | Polysaccharides |
| PWY-6517 | *N*-acetylglucosamine degradation II | Superpathways | Polysaccharides |
| PWY-6558 | heparan sulfate biosynthesis (late stages) | Biosynthesis | Polysaccharides |
| PWY-6566 | chondroitin and dermatan biosynthesis | Biosynthesis | Polysaccharides |
| PWY-6567 | chondroitin sulfate biosynthesis (late stages) | Biosynthesis | Polysaccharides |
| PWY-6568 | dermatan sulfate biosynthesis (late stages) | Biosynthesis | Polysaccharides |
| PWY-6573 | chondroitin sulfate degradation (metazoa) | Degradation/Utilization/Assimilation | Polysaccharides |
| PWY-6576 | dermatan sulfate degradation (metazoa) | Degradation/Utilization/Assimilation | Polysaccharides |
| PWY-66 | GDP-L-fucose biosynthesis I (from GDP-D-mannose) | Biosynthesis | Polysaccharides |
| PWY66-373 | sucrose degradation | Degradation/Utilization/Assimilation | Polysaccharides |
| PWY66-422 | D-galactose degradation V (Leloir pathway) | Degradation/Utilization/Assimilation | Polysaccharides |
| UDPNACETYLGALSYN-PWY | UDP-*N*-acetyl-D-glucosamine biosynthesis<br>II | Biosynthesis | Polysaccharides |
| PWY-6569 | chondroitin sulfate biosynthesis | Superpathways | Polysaccharides |
| PWY-6571 | dermatan sulfate biosynthesis | Superpathways | Polysaccharides |
| PWY-5525 | D-Glucuronate-Degradation | Degradation/Utilization/Assimilation | Polysaccharides |
| PWY-6398 | melatonin degradation I | Degradation/Utilization/Assimilation | SecondaryMetabolism |
| PWY-6399 | melatonin degradation II | Degradation/Utilization/Assimilation | SecondaryMetabolism |
|  | superpathway of geranylgeranyldiphosphate biosynthesis I (via mevalonate) | Superpathways | SecondaryMetabolism |
| PWY-6402 | superpathway of melatonin degradation | Superpathways | SecondaryMetabolism |
| PWY-5120 | geranylgeranyldiphosphate biosynthesis | Biosynthesis | Steroid |
| PWY-5123 | *trans, trans*-farnesyl diphosphate biosynthesis | Biosynthesis | Steroid |
| PWY-5670 | epoxysqualene biosynthesis | Biosynthesis | Steroid |
| PWY-6061 | bile acid biosynthesis, neutral pathway | Biosynthesis | Steroid |
| PWY-6074 | zymosterol biosynthesis | Biosynthesis | Steroid |
| PWY-6132 | lanosterol biosynthesis | Biosynthesis | Steroid |
| PWY-6377 | α-tocopherol degradation | Degradation/Utilization/Assimilation | Steroid |
| PWY66-3 | cholesterol biosynthesis II (via 24,25-dihydrolanosterol) | Superpathways | Steroid |
| PWY66-341 | cholesterol biosynthesis I | Superpathways | Steroid |
| PWY66-377 | pregnenolone biosynthesis | Biosynthesis | Steroid |
| PWY66-378 | androgen biosynthesis | Biosynthesis | Steroid |
| PWY66-380 | estradiol biosynthesis I | Biosynthesis | Steroid |
| PWY66-381 | glucocorticoid biosynthesis | Biosynthesis | Steroid |
| PWY66-382 | mineralocorticoid biosynthesis | Biosynthesis | Steroid |
| PWY66-4 | cholesterol biosynthesis III (via desmosterol) | Superpathways | Steroid |
| PWY-7299 | progesterone biosynthesis | Biosynthesis | Steroid |
| PWY-7306 | estradiol biosynthesis II | Biosynthesis | Steroid |
| PWY-7455 | allopregnanolone biosynthesis | Biosynthesis | Steroid |
| PWY-922 | mevalonate pathway | Biosynthesis | Steroid |
| PWY-7305 | superpathway of steroid hormone biosynthesis | Superpathways | Steroid |
| PWY66-5 | superpathway of cholesterol biosynthesis | Superpathways | Steroid |

## References

Alves R, Chaleil RA, Sternberg MJ. (2002) Evolution of enzymes in metabolism: a network perspective. J Mol Biol. 320(4):751–70. Erratum in: (2002) J Mol Biol 324(2):387.

Bar-Even A, Noor E, Savir Y, Liebermeister W, Davidi D, Tawfik DS, Milo R. (2011) The moderately efficient enzyme: evolutionary and physicochemical trends shaping enzyme parameters. Biochemistry 50(21):4402–10.

Birney E, Clamp M, Durbin R. (2004) GeneWise and Genomewise. Genom Res. 14(5):988–995.

Caspi R, Dreher K, Karp PD. (2013) The challenge of constructing, classifying, and representing metabolic pathways. FEMS Microbiol Lett. 345(2):85–93.

Caspi R, Altman T, Billington R, Dreher K, Foerster H, Fulcher CA, Holland TA, Keseler IM, Kothari A, Kubo A, Krummenacker M, Latendresse M, Mueller LA, Ong Q, Paley S, Subhraveti P, Weaver DS, Weerasinghe D, Zhang P, Karp PD. (2014) The MetaCyc database of metabolic pathways and enzymes and the BioCyc collection of Pathway/Genome Databases. Nucleic Acids Res. 42(Database issue):D459–71.

Duarte, N. C., S. A. Becker, N. Jamshidi, I. Thiele, M. L. Mo, T. D. Vo, R. Srivas, and B. Ø. Palsson. 2007. Global reconstruction of the human metabolic network based on genomic and bibliomic data. Proceedings of the National Academy of Sciences, U.S.A. 104:1777–1782.

Fani R and Fondi M. (2009) Origin and evolution of metabolic pathways. Phys Life Rev. 6(1):23–52.

Grassi L and Tramontano A. (2011) Horizontal and vertical growth of S. cerevisiae metabolic network. BMC Evol Biol. 11:301.

Green ML, Karp PD. (2006) The outcomes of pathway database computations depend on pathway ontology. Nucleic Acids Res. 34(13):3687–97.

Greenberg AJ, Stockwell SR, Clark AG. (2008) Evolutionary constraint and adaptation in the metabolic network of Drosophila. Mol Biol Evol. 25:2537–2546.

Guimerà R, Amaral LAN. (2005). Functional cartography of complex metabolic networks. Nature 433:895–900.

Hudson CM, Conant GC. Expression level, cellular compartment and metabolic network position all influence the average selective constraint on mammalian enzymes. BMC Evol Biol. 2011;11:89.

Innan H, Kondrashov F. (2010) The evolution of gene duplications: classifying and distinguishing between models. Nat Rev Genet 11(2):97–108.

Jensen RA. (1976) Enzyme recruitment in evolution of new function. Annu Rev Microbiol. 30:409–25.

Lazcano A, Díaz-Villagómez E, Mills T, Oró J. (1995) On the levels of enzymatic substrate specificity: implications for the early evolution of metabolic pathways. Adv Space Res. 15(3):345–56.

Light S and Kraulis P. (2004) Network analysis of metabolic enzyme evolution in Escherichia coli. BMC Bioinformatics 5:15.

Light S, Kraulis and P Elofsson A. (2005) Preferential attachment in the evolution of metabolic networks. BMC Genomics 6:159.

Livingstone K, Anderson S. (2009) Patterns of variation in the evolution of carotenoid biosynthetic pathway enzymes of higher plants. J Hered. 100(6):754–61

Lu Y, Rausher MD. (2003) Evolutionary rate variation in anthocyanin pathway genes. Mol Biol Evol. 20:1844–53.

Ma, H., A. Sorokin, A. Mazein, A. Selkov, E. Selkov, O. Demin, and I. Goryanin. 2007. The Edinburgh human metabolic network reconstruction and its functional analysis. Molecular Systems Biology 3:135.

Ma HW, Zeng AP. (2003a) The connectivity structure, giant strong component and centrality of metabolic networks. Bioinformatics 19:1423–1430.

Ma HW, Zeng AP. (2003b) Reconstruction of metabolic networks from genome data and analysis of their global. Bioinformatics 19:270–277.

Milo R, Last RL. (2012) Achieving diversity in the face of constraints: lessons from metabolism. Science 336(6089):1663–7.

Montañez R, Medina MA, Solé RV, Rodríguez-Caso C. (2010) When metabolism meets topology: Reconciling metabolite and reaction networks. BioEssays 32:246–256.

Montanucci L, Laayouni H, Dall’Olio GM, Bertranpetit J. (2011) Molecular evolution and network-level analysis of the N-glycosylation metabolic pathway across primates. Mol Biol Evol. 28(1):813–23.

Nam H, Lewis NE, Lerman JA, Lee DH, Chang RL, Kim D, Palsson BO. (2012) Network context and selection in the evolution to enzyme specificity. Science 337(6098):1101–4.

Noor E, Eden E, Milo R, Alon U. (2010). Central carbon metabolism as a minimal biochemical walk between precursors for biomass and energy. Mol Cell 39(5):809–20.

Notredame C, Higgins DG, Heringa J. (2000) T-Coffee: a novel method for multiple sequence alignments. J Mol Biol 302:205–217.

Peregrin-Alvarez JM, Tsoka S, Ouzounis CA. (2003) The phylogenetic extent of metabolic enzymes and pathways. Genome Res. 13(3):422–7.

Peretó J. (2011) Origin and evolution of metabolisms. In: Origins and Evolution of Life. An Astrobiological Perspective (Gargaud M et al. eds.), Cambridge University Press, ch. 18, p. 270–88.

Peretó J. (2012) Out of fuzzy chemistry: from prebiotic chemistry to metabolic networks. Chem Soc Rev. 41(16):5394–403.

Ramsay H, Rieseberg LH, Ritland K. (2009) The correlation of evolutionary rate with pathway position in plant terpenoid biosynthesis. Mol Biol Evol. 26:1045–1053.

Rausher MD, Miller RE, Tiffin P. (1999) Patterns of evolutionary rate variation among genes of the anthocyanin biosynthetic pathway. Mol Biol Evol. 16:266–274.

Rausher MD, Lu Y, Meyer K. (2008) Variation in constraint versus positive selection as an explanation for evolutionary rate variation among anthocyanin genes. J Mol Evol. 67:137–144.

Romero P, Wagg J, Green ML, Kaiser D, Krummenacker M, and Karp PD. (2004) Computational prediction of human metabolic pathways from the complete human genome Genome Biology 6:R2 R2.1–17.

Storey JD. (2002) A direct approach to false discovery rates. Journal of the Royal Statistical Society, Series B, 64: 479–498.

Thiele, I., N. Swainston, R. M. Fleming, A. Hoppe, S. Sahoo, M. K. Aurich, H. Haraldsdottir, M. L. Mo, O. Rolfsson, and M. D. Stobbe. 2013. A community-driven global reconstruction of human metabolism. Nature Biotechnology 31:419–425.

Teichmann SA, Rison SC, Thornton JM, Riley M, Gough J, Chothia C. (2001) Small-molecule metabolism: an enzyme mosaic. Trends Biotechnol. 19(12):482–6.

Teichmann SA, Rison SC, Thornton JM, Riley M, Gough J, Chothia C. (2001) The evolution and structural anatomy of the small molecule metabolic pathways in Escherichia coli. J Mol Biol. 311(4):693–708.

Vitkup D, Kharchenko P and Wagner A. (2006) Influence of metabolic network structure and function on enzyme evolution. Genome Biol. 7:R39.

Yang Z. (2007) PAML 4: phylogenetic analysis by maximum likelihood. Mol Biol Evol 24(8):1586–1591.

Yčas, M. (1974). On earlier states of the biochemical system. J Theor Biol 44:145–60.

Zhao J, Yu H, Luo JH, Cao ZW, Li YX. (2006) Hierarchical modularity of nested bow-ties in metabolic networks. BMC Bioinformatics 7:386.

